# 2-kupl: mapping-free variant detection from DNA-seq data of matched samples

**DOI:** 10.1101/2021.01.17.427048

**Authors:** Yunfeng Wang, Haoliang Xue, Christine Pourcel, Yang Du, Daniel Gautheret

## Abstract

The detection of genome variants, including point mutations, indels and structural variants, is a fundamental and challenging computational problem. We address here the problem of variant detection between two deep-sequencing (DNA-seq) samples, such as two human samples from an individual patient, or two samples from distinct bacterial strains. The preferred strategy in such a case is to align each sample to a common reference genome, collect all variants and compare these variants between samples. Such mapping-based protocols have several limitations. DNA sequences with large indels, aggregated mutations and structural variants are hard to map to the reference. Furthermore, DNA sequences cannot be mapped reliably to genomic low complexity regions and repeats. Herein, we introduce 2-kupl, a k-mer based, mapping-free protocol to detect variants between two DNA-seq samples. On simulated and actual data, 2-kupl achieves a higher precision than other mapping-free protocols. Applying 2-kupl to prostate cancer whole exome data, we identify a number of candidate variants in hard-to-map regions and propose potential novel recurrent variants in this disease.

## Introduction

Searching for genomic variants is a fundamental aspect of medical research, whether in the study of Mendelian diseases or of somatic, cancer-related alterations[1]. While certain variants result in gene dysfunction and disease[2], others are largely asymptomatic but give rise to neoantigens relevant to immune escape and therapeutic efficacy or treatment[3]. Genome variants are also of interest in microbiology to analyze the differences between microbial strains[4] and reveal mechanisms underlying phenotypes. In this study, we address the problem of finding genomic differences between a matching pair of high throughput DNA sequencing (DNA-seq) datasets from the same individual (human somatic variation) or from two bacterial strains.

Genomic variants include mutations, indels and structural variants (SV). Mutations and indels can alter genes by disrupting the genetic code, while SVs, by pulling distant regions together or splitting one region into segments, can create chimeric genes or have a broader impact on whole chromosomal regions[5]. Variants are typically detected by whole-genome (WGS) or whole-exome (WES) sequencing through comparison with reference sequences. Aligners such as BWA[6] are first applied to map reads to the reference sequences. The variant calling step then detects differences between mapped reads and the reference. Popular variant callers include MuTect2[7], Varscan[8], somaticsniper[9], MuSE[10]. Based on variants observed between two sequence samples and a common reference genome, these programs can then infer differences between the two samples (e.g., in Mutect2’s somatic mode).

Reference-based variant calling has well-known limitations. Aligners may encounter difficulties while handling reads with low mapping qualities[11], originating from repeat regions, low complexity regions or complex variants. These reads of low mapping quality are usually discarded. Furthermore, some species have no reliable reference, which is common in microbes[12].

Alternative approaches to variant calling involve mapping-free protocols[13]. These methods do not rely on a reference genome and can directly predict variants from the raw fastq file. A typical strategy is to use a de Bruijn graph (DBG)[14]. A DBG is constructed using *k*-mers (subsequences of fixed size *k*) decomposed from the sequence reads. The occurrence of *k*-mers harboring a mutant allele and a wild type allele generates a bubble structure in the DBG. Variant callers developed based on DBGs include DiscoSNP++[15] and Lancet[16]. DBG-based methods also introduce new issues. First, complex genomic variants and repeats may result in complicated graphs that are difficult to parse[17]. Second, short contigs may be discarded at the post-processing step, where branch pruning may cause many false negatives. Furthermore, sequences assembled by *k*-mers without variants have little contribution if the purpose is detecting variants. Only reconstructing the active regions spanning the variants is more efficient than considering all kmers[13]. Although it is possible to extend DBG-based methods to SV detection, the lack of sensitivity to local events makes these approaches less suitable for finding variants in ambiguous regions, such as repeats[18]. This motivates the need for a method to detect variants in arbitrary genome regions directly from DNA-seq data.

We present 2-kupl, a *k*-mer-based bioinformatic pipeline that compares matched case and control samples to discover case-specific variants. 2-kupl extracts case-specific *k*-mers and the matching counterpart k-mers corresponding to a putative mutant and reference sequences and merges them into contigs. As 2-kupl begins with a shortlist of cs-kmers, the number of *k*-mers considered from unaltered regions and non-specific variants is drastically reduced compared with DBG-based methods.

We compare the accuracy and CPU-requirements of 2-kupl, DiscoSNP++, Lancet and GATK-Mutect2, using both simulated and real DNA-seq datasets. We analyze the nature of novel variants detected by 2-kupl and potential reasons for their absence in conventional protocols. We also use 2-kupl to detect recurrent variants in prostate cancer WES samples from the TCGA project[19]. Finally, we evaluate 2-kupl precision in bacterial WGS data. Overall, we demonstrate that 2-kupl is a practical and powerful alternative to complement the discovery of genomic variants in hard-to map regions or species with no reliable reference.

## Methods

### Outline of 2-kupl pipeline

The general pipeline is presented in Fig. 1. The input is composed of DNA-seq data from two matched samples. Samples typically correspond to control/normal/wild-type and a case/tumor/mutant-type. For cancer data, we strongly recommend using as a control of a distant tissue such as white blood cells rather than adjacent normal tissues, as the later can be contaminated by tumor cells and 2-kupl only considers variant sequences that are absent in the control dataset. Sequence types can be either single-end or paired-end sequencing reads. 2-kupl then identifies pairs of case-specific k-mers (cs-kmers) and counterpart k-mers (ct-kmers). 2-kupl returns predicted variants exclusive to the case sample, including mutations, indels and structural variation. Variant statistics including cs-count, coverage, allele frequency and variant P-value are computed. A variant file and an alignment file are produced.

**Figure 1.**
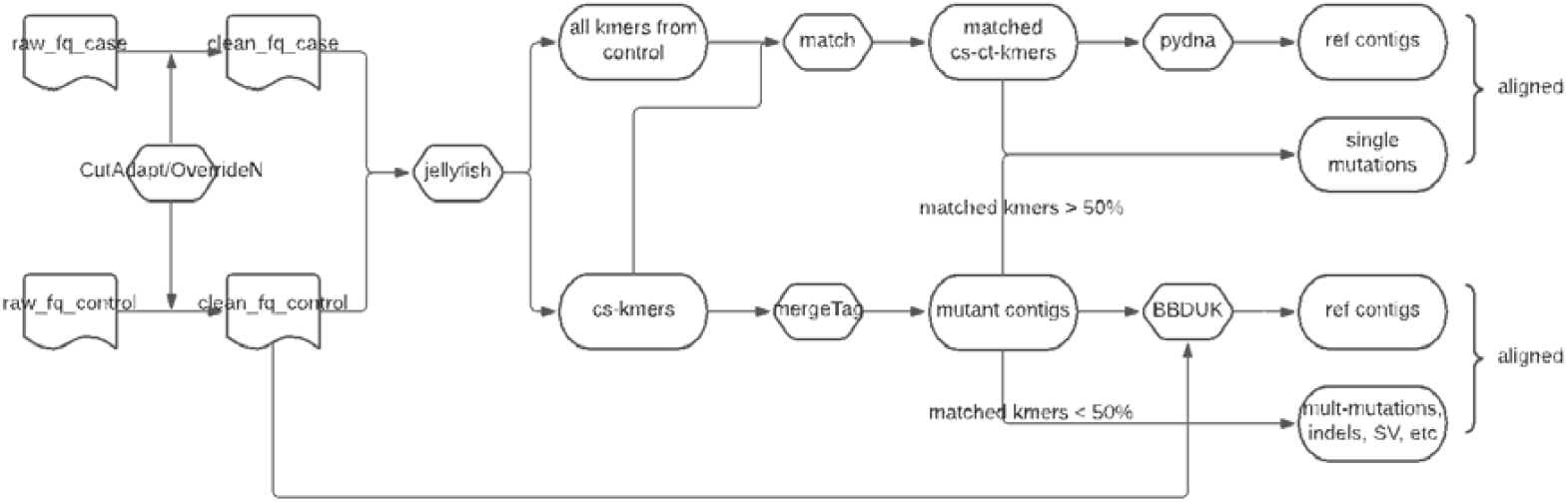
Overall workflow of 2-kupl.

2-kupl is developed purely in Python. The main dependencies include Jellyfish[20] and GSNAP[21]. Other dependent python libraries and instructions can be found from the Github repository https://github.com/yunfengwang0317/2-kupl

### Data cleaning

Low quality sequences are trimmed with Cutadapt[22] (parameter ‘--quality-cutoff’ = 10). As CutAdapt does not remove low-quality bases within the central part of reads, we implemented an overriding function that replaces each low-quality base (Phred score <10) with N. This procedure is applied to both case and control libraries.

### k-mer indexing and counting

Jellyfish is used to index and quantify k-mers from both case and control with options k=31 and −C (canonical k-mers). As Jellyfish removes k-mers containing Ns, none of the low-quality bases is present in the k-mer list. The generated k-mers subsequently undergo two filtering steps. First, k-mers with counts below a user-specified cutoff (default=3) are removed. These low abundance k-mers are assumed to result from sequencing errors or off-target regions in the case of WES data. Second, k-mer lists from case and control are compared and only case-specific k-mers (cs-kmers) are retained.

### Matching counterparts of cs-kmers

For each cs-kmer harboring a point mutation, there should exist a counterpart k-mer (ct-kmer) from the control dataset with only one base substitution (Hamming distance =1), which can be considered as a product of the wild type sequence. Note that Hamming distance=1 only considers substitutions. Hence single nucleotide insertions and deletions are rejected at this step and will be treated later with unmatched k-mers. Finding the matched ct-kmer for each cs-kmer should allow us to infer the variation without reference sequences. We initially build a hash table where the keys are the continuous 15 bases from each side of cs-kmers. For each 15-bases key, we create a bucket of all k-mers starting or ending with the key. Then we survey the buckets and seek all k-mer pairs with a hamming distance of 1 in the same bucket. We thus generate all k-mer pairs (k_i_,k_j_) with a Hamming distance of one. For any pair of k-mers with a Hamming distance of 1, if one k-mer comes from the cs-kmer list and the other comes from the control, this pair of k-mers is considered to be matched. Otherwise, we allocate the cs-kmers to the “unmatched k-mers” group. These unmatched k-mers either contain variants of more than one nucleotide (multiple mutations, indels and structural variants) or come from low coverage regions. The schematic workflow is shown in Fig. 2.

**Figure 2.**
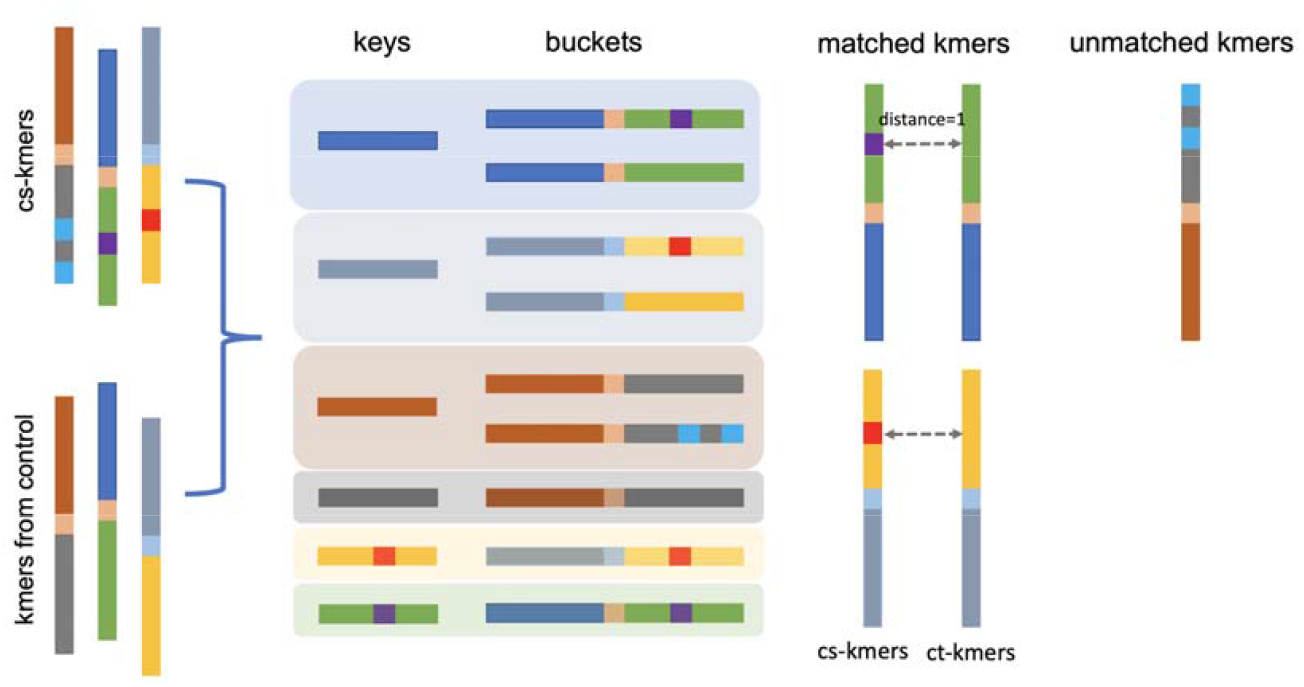
Procedure for matching cs-kmers to ct-kmers. Long rectangles represent one 31-mer. Short rectangles (keys) represent the head or tail 15 bp of a cs-kmer. Color changes indicate sequence differences.

### Assembly of cs-kmers into mutant contigs

cs-kmers are assembled into mutant contigs that correspond to variants and their local context. The assembly process is done using the “mergeTag” function from DEkupl[23] (https://github.com/Transipedia/dekupl). Two k-mers overlapping by k-i bases are merged iteratively with i ranging from 30 to 25 (min_overlap parameter is set to 25 by default). The merging process is interrupted when no k-mers can be added or ambiguity occurs (two different overlapping k-mers are encountered).

### Inferring reference contigs

We use two distinct procedures for reference sequence determination, depending on whether or not sufficient ct-kmers are available to build a reference contig. For each mutant contig, if more than half of its component k-mers are matched, all the ct-kmers are merged by the python package pydna[24]. The resulting mutant contigs correspond to isolated mutations. Merged contigs produced by ct-kmers can be regarded as putative references. For each pair of mutant and reference contig, we then define two values representing counts of supporting k-mers for the mutant allele (cs-count) and supporting k-mers for both mutant and reference alleles (coverage). The cs-count is computed from the median k-mer count of cs-kmers and coverage is calculated from the sum of the median count of cs-kmers and ct-kmers. Herein, we select the median count instead of the mean count because mean values are more sensitive to high-count k-mers from repeats or copy number amplification regions.

For mutant contigs in which less than half of the k-mers are paired, we consider that a reference cannot be assembled from paired-kmers. A procedure was implemented to retrieve the reference from the original reads. Reads with at most one mismatch to any k-mer from the mutant contig are retrieved from the control fastq file using BBDUK[25]. These reads are then assembled by CAP3[26]. In this way, we can infer the putative reference for each contig and evaluate coverage based on the number of reads retrieved by BBDUK. The cs-kmers in these contigs have no matching ct-kmers and contigs are thus considered to contain multiple mutations, indels and structural variants.

### Filtering low-quality variants

The cs-count and coverage substantially impact the reliability of events called by 2-kupl. For instance, a sequencing error could be repeatedly generated in a region of high coverage. Besides, sequencing errors may, by chance, be detected as mutations with high allele frequency in low coverage regions. Thus, false positives are introduced due to either high cs-count in high coverage regions or high allele frequency in low coverage regions. However, coverage varies between whole-genome sequencing (WGS) and whole-exome sequencing (WES) data. WGS does not use an upfront enrichment step so it generates a more uniform coverage of the genome. On the other hand, the enrichment steps involved in WES lead to non-uniform coverage, generating coverage ‘hot’ and ‘cold’ spots[27].

2-kupl provides several criteria for users to evaluate call reliability. A Fisher’s exact test P-value is calculated based on the cs-count and coverage in tumor and matched normal libraries for each variation. A Phred quality score is subsequently computed as-10*log*_l0_*P*. Users can specify cutoffs for cs-count, coverage, allele frequency and Phred to filter results. Default cutoffs for cs-count, coverage, allele frequency and Phred are set to 3, 10, 0.05 and 5, respectively.

### VCF format export

Events identified by 2-kupl are exported as a variant call format (VCF) file[28]. 2-kupl outputs the contig harboring the variation and the corresponding putative reference without the variation for each event. If users provide an available reference, the mutant contig is mapped to this reference using GSNAP[21]. After the mapping process, actual chromosome and position information are provided in the VCF file. Besides the VCF file, 2-kupl also exports an alignment of each contig and its putative reference obtained using the pairwise2 python package[29]. Contigs corresponding to indels and structural variants are further mapped to reference by BLAST[30] (default parameters) which we found better suited to fragmented alignments.

### Comparison with other software

We compared 2-kupl with three other tools. DiscoSNP++[15] is designed for detecting SNVs and small indels from fastq files without using reference. DiscoSNP++ first decomposes the reads into k-mers and constructs a DBG[31]. The context contigs can be extracted from DBG bubbles that correspond to local variants. Lancet[16] relies on localized colored DBG to detect somatic variants in paired samples. K-mers shared by two matched samples or specific to either of them are marked in different colors in the DBG. In this way, Lancet is able to detect case-specific events. It is worth mentioning that Lancet uses bam format files as input so it also leverages the reference before variant detection. We also compared 2-kupl with the leading reference-based GATK-Mutect2 pipeline[7]. GATK-Mutect2 takes mapped sequence files as input, detects variants based on the reference and compares the variants of two matched samples to identify case-specific variants (somatic mode). Version hg38 of the human genome was used in all reference-based procedures. To make runtime comparisons fair, we took the mapping procedure into account in Lancet and GATK-Mutect2. Alignment was performed using BWA with default parameters. Thus all four protocols started with fastq files. To evaluate the dependency of 2-kupl running time on the number of k-mers, we ignored the part up to k-mer counting. Mapped reads were visualized with the Integrative Genomics Viewer (IGV)[32] 2.6.2 on hg38.

### Simulated WES analysis

We downloaded simulated WES data from Meng and Chen[33]. This dataset was developed based on the NA12878 pilot genome[34] (reference data set of 5.4 million phased human variants validated by genetic inheritance from sequencing a three-generation 17-member pedigree). The authors used BAM-Surgeon[35] to select genomic loci and introduce random SNV and indel spike-ins, and generated 2×100nt reads WES files at 230X coverage. For our benchmark, we used a tumor sample described by authors as one of the most complicated, NA12878_79_snv_indel_sorted.bam (with four sub-populations, expected VAFs of 0.5, 0.35, 0.2 and 0.1). Picard was used to convert bam files to fastq format files with default parameters. 2-kupl was run using default parameters on pairs of simulated normal-tumor fastq files.

### TCGA-PRAD data analysis

Matched normal-tumor WES data of 498 patients from TCGA-PRAD[36] were retrieved with permission from dbGAP[37]. BAM files were converted to paired-ends fastq files using Picard tools with default parameters. 2-kupl somatic variant calls were obtained for each normal/tumor pair using default parameters. Detailed analysis of variant calling was performed on the TCGA PRAD sample with the highest tumor mutational burden (barcode TCGA-ZG-A9ND).

2-kupl results on the TCGA-PRAD dataset were compared to variant calls downloaded from the GDC portal. Briefly, the GDC portal workflow uses BWA to map reads to the human genome and determines variants with five state of the art variant callers. We used the maftools R package[38] to retrieve variants predicted using the GATK-Mutect2 pipeline and filtered against a “panel of normals”. This mutation dataset is hereafter referred to as the “GDC portal” dataset. GDC portal pipelines are detailed at: https://docs.gdc.cancer.gov/Data/Bioinformatics_Pipelines/DNA_Seq_Variant_Calling_Pipeline/.

To remove putative germline variants from 2-kupl results, we built a boolean matrix representing the presence of each k-mer in each normal sample. Any k-mer present in at least two normal samples was excluded. Retained recurrent variants were considered as tumor-specific (Suppl. Table S1). Mutations detected by 2-kupl and absent in the GDC portal variants were considered as 2-kupl specific. To verify whether calls absent in GDC portal variants were not discarded at earlier stages of the GDC portal pipeline, we also retrieved the protected MAF file containing all unfiltered variants called by the Mutect2 workflow.

The oncoplot graph for GDC portal variants (Fig. 7A) was drawn using maftools. To obtain recurrently mutated genes by 2-kupl, we aggregated variants belonging to the same gene in 2-kupl results and constructed a gene-level occurrence matrix that was fed to maftools (Fig. 7B). Recurrent variants from 2-kupl and the GDC Portal were also compared with a comprehensive prostate cancer dataset from 200 whole-genome sequences and 277 whole-exome sequences from localized prostate tumours[36] (Suppl. Table S2)

Recurrently mutated genes were annotated using a collection of 1404 PRAD-related genes collected from CLINVAR[39], COSMIC[40], DISEASE[41], KEGG[42], OMIM[43], PheGenl[44] and driver predictions by Martincorena *et al*. and Armenia *et al*. [45, 46] (Suppl. Table S3).

### Bacterial genome analysis

We obtained WGS fastq files from the *Pseudomonas aeruginosa* PAO1Or wild-type strain and 24 phage-tolerant mutants[47]. Mutations in the phage-tolerant variants were previously validated by mapping of the WGS raw sequences to the PAO1Or genome (Genbank accession LN871187) and confirmed by PCR amplification and Sanger sequencing. We used one control WGS file and 21 mutant WGS files corresponding to 26 validated variants. Detailed variants (Suppl. Table S4) include seven mutations, 13 small indels and six large deletions longer than 100 bp. 2-kupl was run using default parameters on every mutant WGS file compared to the control WGS file.

## Results

### A novel algorithm for detecting variants between two DNAseq samples

We developed 2-kupl to predict variants between pairs of matched DNAseq libraries. Input libraries consist of a “mutant type” and a “wild type” sample such as a pair of tumor and normal tissues from one patient or a pair of mutant and wild-type bacterial cells. Data can be either WGS or WES. 2-kupl identifies sequence fragments (contigs) specific to the mutant dataset and their wild-type counterpart in the control dataset. This operation is done without relying on a reference genome. If a reference genome is provided, 2-kupl can also align contigs to the reference and generate genomic coordinates just like with mapping-based methods.

### Performance on simulated WES data

We first applied 2-kupl to the detection of somatic mutations in a simulated human cancer WES dataset containing a known number of spliked-in mutations. We compared 2-kupl with three other software, including two mapping-free methods (DiscoSNP++ and Lancet) and the leading mapping-based pipeline GATK-Mutect2. Results are summarized in the first column of Table 1. The number of cs_kmers to process is reduced by nearly 20% after data cleaning by 2-kupl. 88.6% of cs_kmers were matched to ct_kmers, corresponding to predicted point mutations or indels. We evaluated mutations and indel calls by 2-kupl and concurrent methods (Table 2). For mutation calling, 2-kupl performed better than the other mapping-free methods in terms of false-positive, FDR, precision and F1 score (Table 2). Lancet and GATK achieved better recall than 2-kupl, but Lancet also introduced more false positives. 2-kupl had a higher recall for calling indels than DiscoSNP++ and Lancet but was outperformed by DiscoSNP++ in FDR and Precision (Table 3). Expectedly, GATK-Mutect2 outperformed all mapping-free approaches regardless of variant types. DiscoSNP++ did not perform as well as others in this comparison because it calls variants in a single sample. To achieve the comparison between two matched samples, we ran DiscoSNP++ on each sample separately and subsequently removed all variants that occurred in both samples. Lancet performed well in terms of recall but at a high cost of false positives. As expected, most false positives had few reads containing the alternative allele, which is frequent with Lancet. The high recall and high rate of false positives produced by Lancet are consistent with the conclusions of Meng and Chen[33]. The GATK-Mutect2 pipeline outperformed all mapping-free approaches when calling mutations. The use of a reference sequence and the Haplotype Caller algorithm gives GATK-Mutect2 a clear advantage. Even though 2-kupl got a relatively lower recall than GATK-Mutect2, it had better control of the false positives and got a higher precision when calling indels (Table 3).

**Table 1.** Number of k-mers and contigs after applying 2-kupl on two matched libraries

|  | simulated WES | TCGA-ZG-A9ND WES |
| --- | --- | --- |
| all k-mers(Tumor/Normal) | 465,718,268/465,610,133 | 184,233,006/177,517,776 |
| raw cs_kmers | 23599 | 393525 |
| cleaned cs_kmers | 18439 | 291350 |
| matched cs_kmers | 16914 | 240360 |
| all contigs | 1245 | 106426 |
| mutations | 1026 | 9901 |
| indels | 112 | 1105 |
| unmapped | 0 | 58 |
| low confidence | 107 | 312 |

**Table 2.** Comparison of four approaches on mutations using simulated WES data

| mutation | 2-kupl | DiscoSNP++ | Lancet | GATK_MuTect2 |
| --- | --- | --- | --- | --- |
| True Positive | 581 | 107 | 604 | 689 |
| False Positive | 45 | 85 | 126 | 2 |
| False Negative | 241 | 715 | 218 | 133 |
| Recall | 0.71 | 0.13 | 0.73 | 0.84 |
| FDR | 0.07 | 0.44 | 0.17 | 0.003 |
| Precision | 0.93 | 0.56 | 0.83 | 0.997 |
| F1 score | 0.80 | 0.21 | 0.78 | 0.91 |

**Table 3.** Comparison of four approaches on indels using simulated WES data

| indels | 2-kupl | DiscoSNP++ | Lancet | GATK_MuTect2 |
| --- | --- | --- | --- | --- |
| True Positive | 42 | 14 | 40 | 49 |
| False Positive | 16 | 2 | 44 | 26 |
| False Negative | 39 | 67 | 41 | 32 |
| Recall | 0.52 | 0.17 | 0.49 | 0.60 |
| FDR | 0.27 | 0.13 | 0.52 | 0.35 |
| Precision | 0.72 | 0.88 | 0.47 | 0.65 |
| F1 score | 0.60 | 0.29 | 0.48 | 0.63 |

Another advantage of 2-kupl is the short running time (Fig. 3A). 2-kupl took 1.6 hours to analyze the simulated WES data with default parameters. DiscoSNP++ took 2.54 hours to run on both case and control samples in parallel. Both Lancet and GATK-Mutect2 require prior mapping of reads to the human genome (which takes 3.17 hours), explaining in part their longer runtimes.

**Figure 3.**
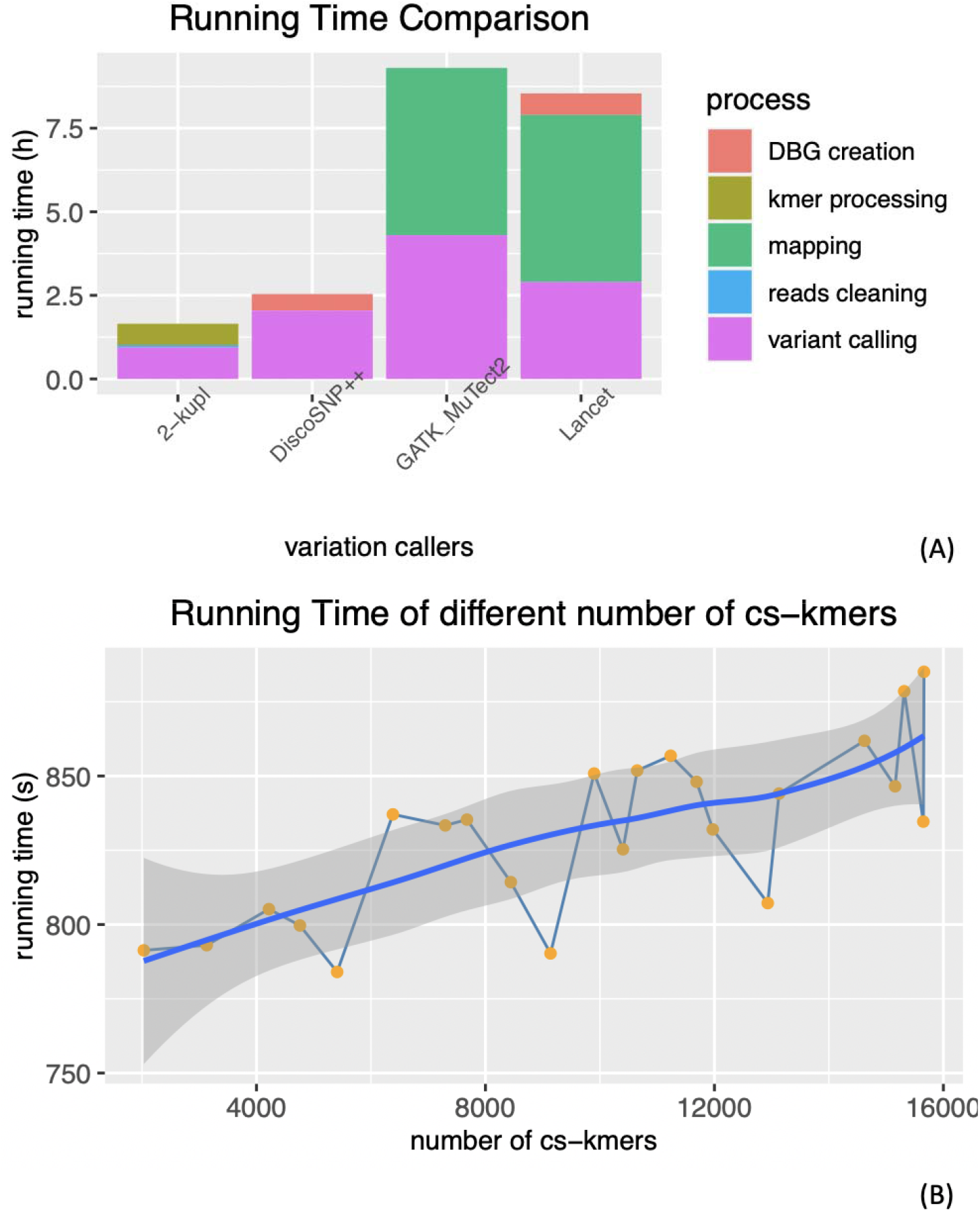
Running time and performance with different types of variants (A) Overall running times of four softwares. (B) Running times of 2-kupl for different numbers of k-mers.

To evaluate 2-kupl run time dependency on the number of cs_kmers, we ran 2-kupl on datasets with different numbers of cs_kmers (Fig. 3B). Running time increased linearly with the number of cs-kmers. Each additional 10,000 cs-kmers increased the running time by nearly 50 seconds.

We estimated the performance of 2-kupl under different parameter combinations. Coverage and cs-count thresholds (‘min_cov’ and ‘min_cs-count’, respectively) were varied from 3 to 9. Results are shown in Fig. 4. The min_cs-count parameter was negatively related to recall and positively related to false negatives. The min_cov parameter was inversely related to recall, FDR, recall and false positives. Precision reached an inflection point when min_cscount was set to 4.

**Figure 4.**
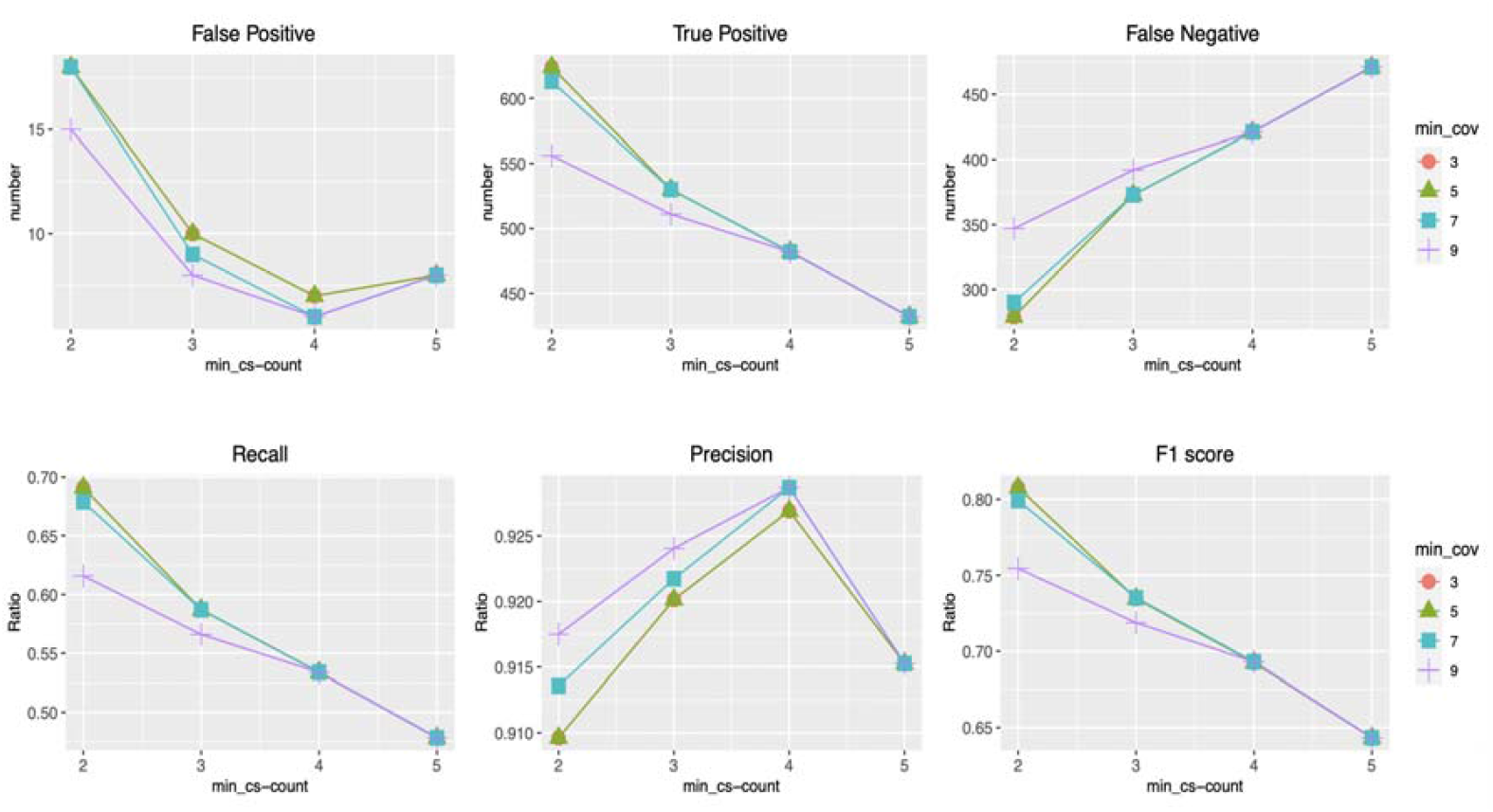
Robustness of 2-kupl using different parameters. The x-axis indicates the min_cs-count parameter and the y-axis represents the corresponding ratio or number. The trend lines under different min_cov parameters are represented by four colors.

### Assessing 2-kupl on a real normal-tumor WES dataset

To assess 2-kupl results on actual WES data, we applied 2-kupl on one WES dataset of matched tumor and normal tissues from the TCGA-PRAD dataset. We firstly compared 2-kupl and GDC portal somatic variant calls (obtained with GATK-Mutect2) on the TCGA patient with the highest tumor mutational burden. The numbers of k-mers, contigs and variants obtained by 2-kupl are shown in Table 1. Mutation calls by 2-kupl and GDC portal variants are shown in Table 5. Although total call numbers were similar, only 327 calls (∼9%) were shared by the two approaches, including 319 mutations and 8 indels. Among the variants detected by 2-kupl, 193 (5.13%) mapped to noncoding regions and 101 (2.7%) were annotated as repeats by RepeatMasker[48]. 2-kupl also captured 57 (1.5%) unmapped variants. 173 2-kupl variants (4.6%) were mapped to low mappability “blacklist” regions[49]. In spite of the small general overlap of 2-kupl and GDC portal variants, the two methods show significantly higher agreement on high scoring 2-kupl calls (Fig. S1).

**Table 5.** Number of mutations and indels detected by 2-kupl and GDC portal variants

|  | 2-kupl | GDC portal variants | overlap |
| --- | --- | --- | --- |
| mutation | 3607 | 3093 | 319 |
| indel | 151 | 823 | 8 |
| total | 3758 | 3916 | 327 |

We further analyzed mutations specific to 2-kupl. These calls may have been rejected in GDC portal variants for a number of valid reasons, including low mapping quality, location in short tandem repeats or presence in normal samples. A real “miss” by the reference-based pipeline should be recorded only when reads could not possibly be aligned to the genome while they indeed contained a valid mutation.

Fig. 5A shows a case of false positives introduced due to artifactual cs-kmers. Generally, k-mers harboring a mutation present in both tumor and normal tissues are supposed to be ruled out. However, erroneous tumor-specific “cs_kmers” can escape the filtering process if the same k-mer in the normal tissue happens to be low quality and is discarded.

**Figure 5.**
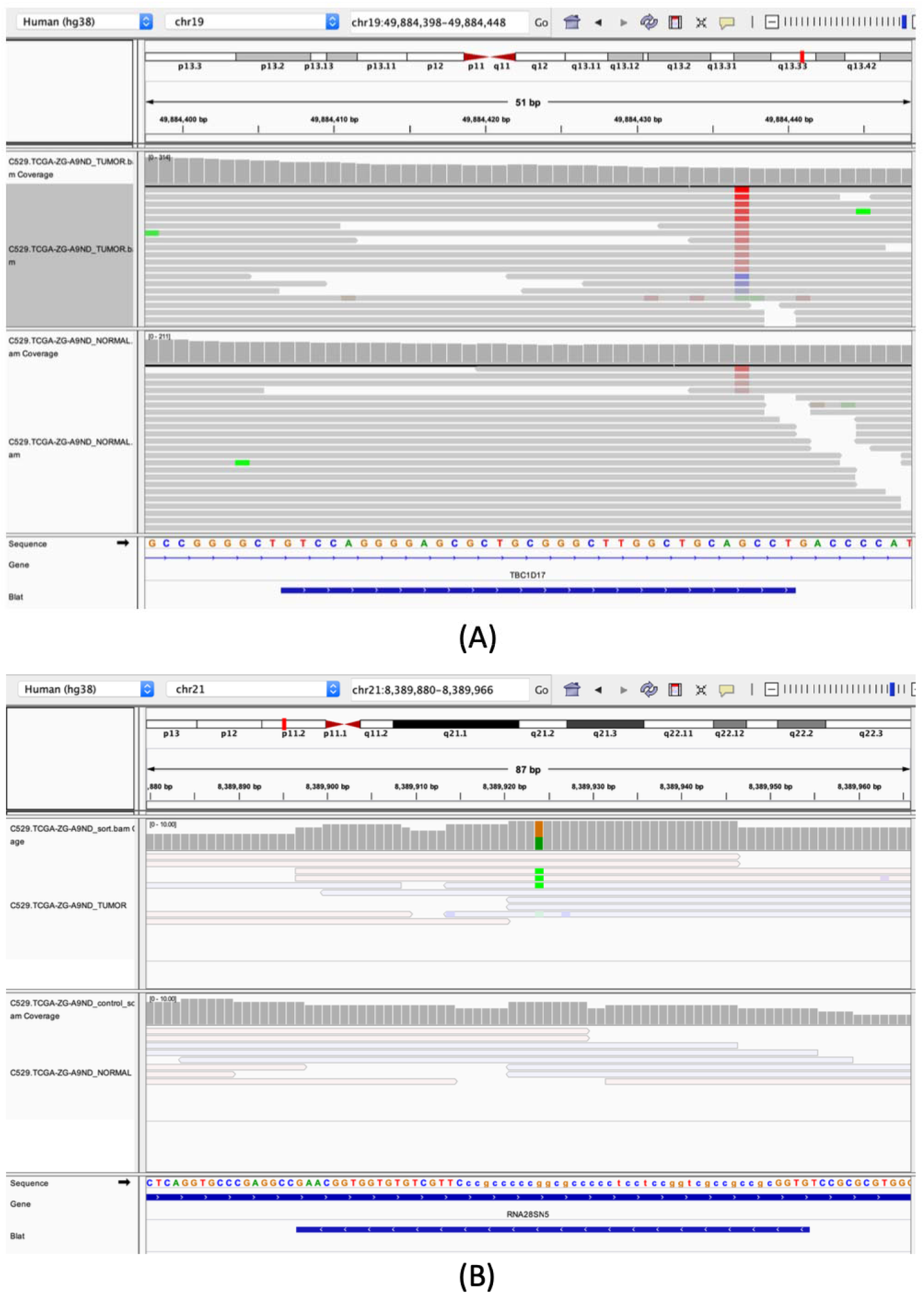
IGV views of variant calls in TCGA-PRAD WES dataset. The two central tracks show aligned reads from the tumor (top) and normal (bottom) WES library. The lower track shows gene annotation and 2-kupl contigs. (A) A likely false-positive call by 2-kupl at a position of low mapping quality (B) A likely true positive within a repeat region. Reads in transparent color have low MAPQ values (<10).

Certain 2-kupl specific mutations are possibly true positives discarded by mapping-based protocols due to their location within a repeat region. Fig. 5B shows such a potential somatic mutation. The mutation is located within a ribosomal RNA gene that is repeated multiple times in the genome and further contains a C-rich repeat (represented in lower cases). Reads generated from these repetitive regions are given low MAPQ values by mappers and variants in these regions are then discarded by variant callers.

Among the unmapped 2-kupl calls, only one has a Phred score in the top 5% (Fig. S2). The mutant sequence and its inferred reference are shown in Fig. S3. The mutant contig is covered by 0 and 47 reads in the Normal and Tumor sample, respectively while the reference is covered by 88 and 65 reads in the Normal and Tumor sample, respectively (Fig. 6). The sequence maps to a centromeric repeat of Chr22, with three mismatches. The mapping procedure would thus miss this highly significant variant.

**Figure 6.**
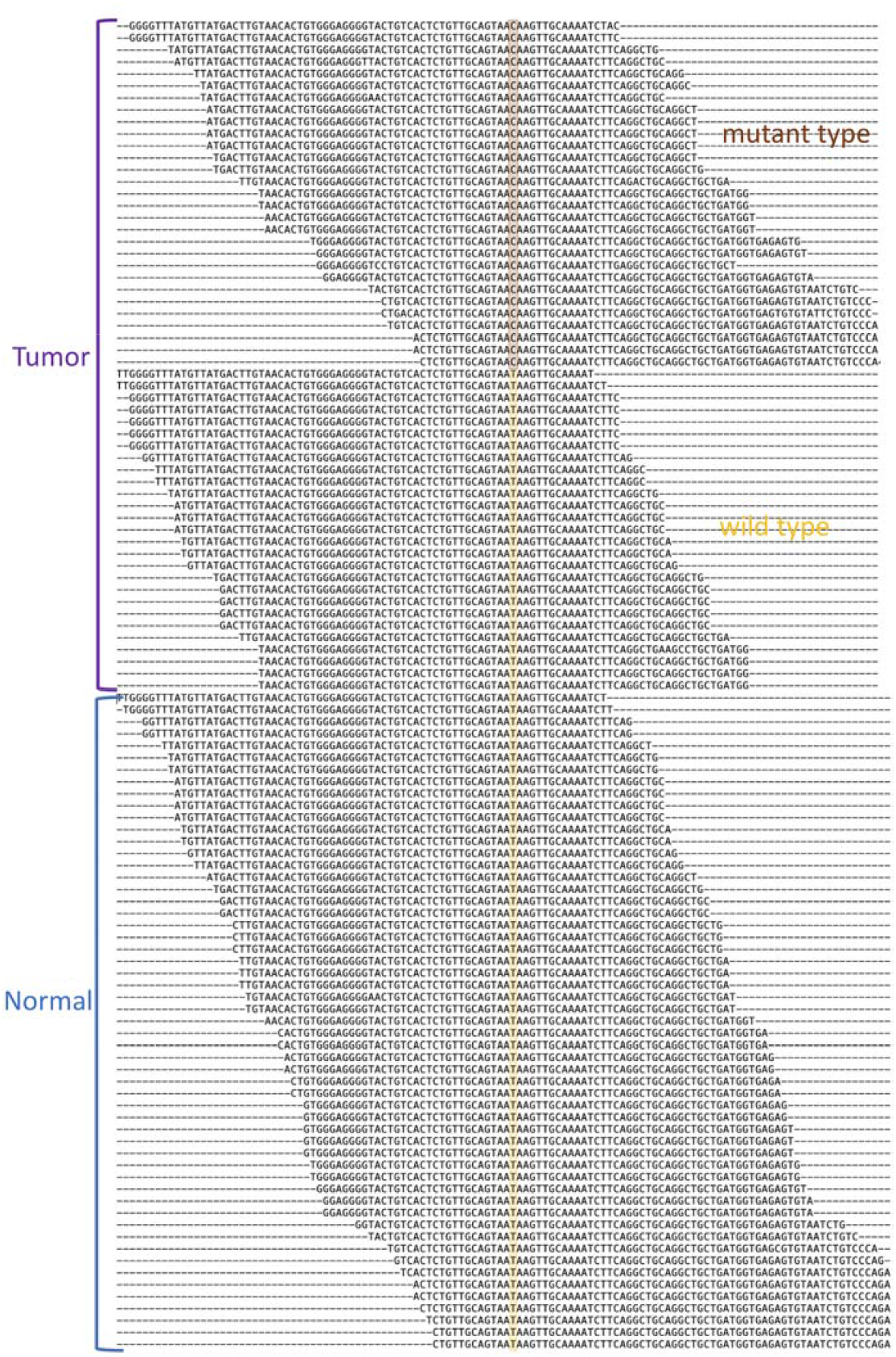
An unmapped somatic variant from a TCGA PRAD patient. Only reads matching the central k-mer of the tumor-specific variant or its inferred counterpart are shown. Reads from the tumor and normal samples are distinguished. The position of variation is highlighted.

### Recurrent mutations in TCGA-PRAD

Recurrence across patients is a powerful criterion for distinguishing drivers from passenger mutations[50–52] and has been used to discover drivers and define molecular subtypes of prostate cancer[53]. We applied 2-kupl to each pair of Normal/Tumor samples in the complete PRAD WES dataset (N=248) and identified 3211 recurrent variants (suppl. Table S1). For comparison we retrieved from the GDC portal recurrent variants predicted for the same dataset (GATK-Mutect2 pipeline, see Methods). Among 3734 recurrent variants in the GDC portal, 854 were shared with 2-kupl recurrent variants (suppl. Table S1). We further compared the recurrent variants to a comprehensive dataset of recurrent prostate cancer mutations from Fraser *et al*.[54] based on 200 whole-genome and 277 whole-exome sequences from multiple sources. Comparisons were restricted to exonic regions. Over 48 recurrent mutations in Fraser *et al*, a similar number was shared with 2-kupl or the GDC-portal (22 and 21, respectively) (Suppl. Table S2). Among recurrent mutations specific to 2-kupl, we note the one found at chr14:37592023 within an exon of *FOXA1*, a putative prostate cancer driver [45], in 3 TCGA-PRAD patients.

We further compared 2-kupl calls to GDC portal variants at the level of genes (Detailed in Method section). The GDC portal reported 6944 genes mutated in 2 or more patients, vs. 14137 recurrent genes by 2-kupl. Enrichment analysis shows a good convergence of the most frequently mutated genes by the two methods (Fig. 7A). Fig. 7B and 7C show oncoplot views of the top 20 genes according to the GDC portal and 2-kupl, respectively, showing eight shared genes. Both gene lists are contaminated by long (*TTN*) or highly polymorphic genes (Mucins) whose recurrence is an artifact due to higher mutation counts. Although software is available to account for those effects[55], we purposely analyze the uncorrected list of genes here. Among the top 20 mutated genes by 2-kupl and GDC portal, 7 and 9 genes, respectively, are known prostate cancer-related genes. Among those, *UBR4, DNAH5* and *LRP1* were only detected by 2-kupl. When considering the top 50 recurrently mutated genes according to 2-kupl and GDC portal, 19 and 23, respectively, are cancer-related. Among those, *HSPG2, DNAH3, UBR4, COL6A3, CABIN1, IGF2R, PTPRF, DNAH5, HTT* and *TRRAP* were only detected by 2-kupl.

**Figure 7.**
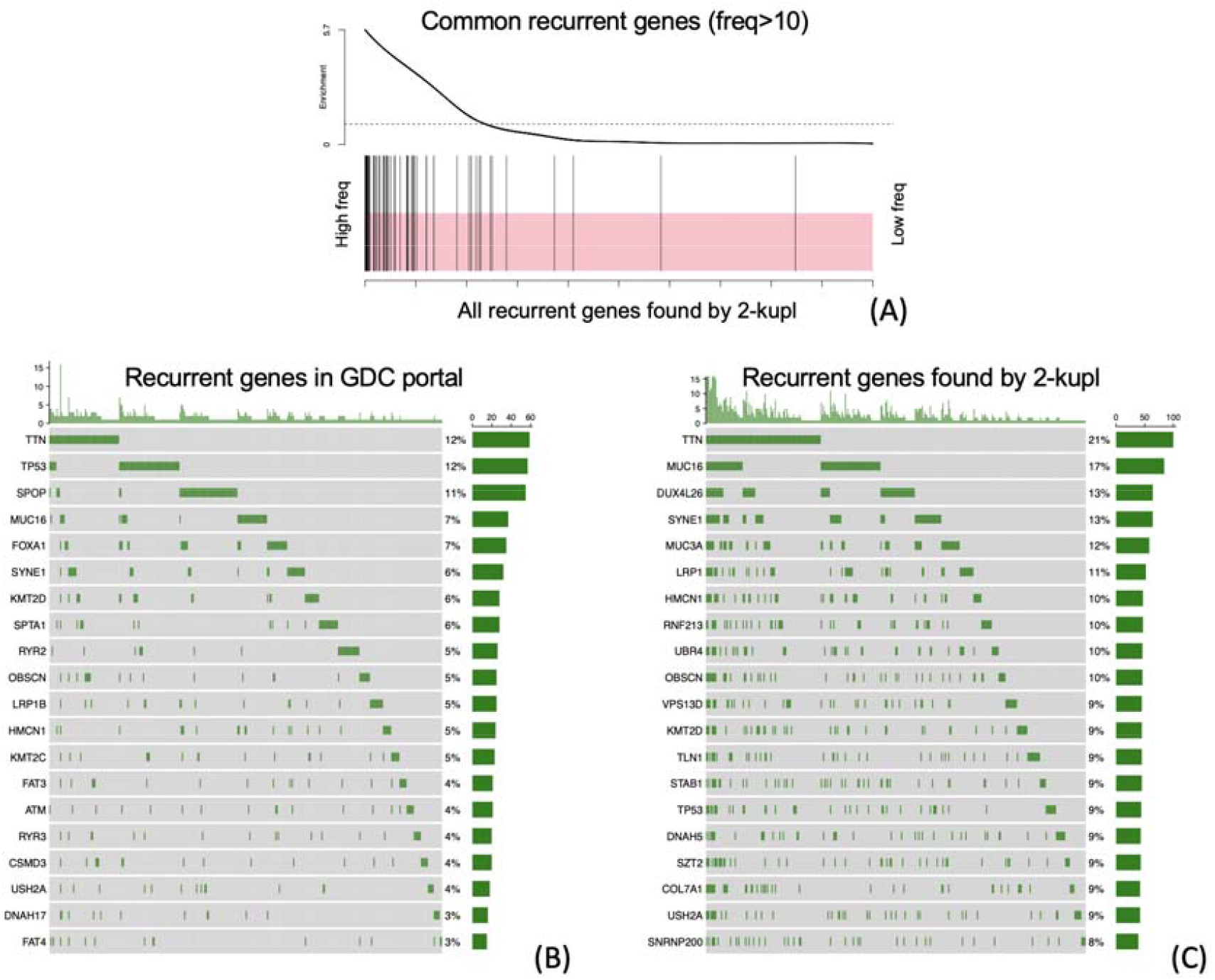
Recurrently mutated genes in the TCGA-PRAD WES dataset. (A) Enrichment analysis of recurrent genes. The vertical bars are the common recurrently mutated genes (altered in at least ten patients) between GDC portal and 2-kupl. The x axis represents the recurrent genes found by 2-kupl sorted by frequency. The smooth curve reflects the degree to which the common genes are overrepresented in the whole 2-kupl recurrent genes. (B) The 20 genes with the highest mutational frequency detected in GDC portal variants. (C) The top 20 recurrent genes with the highest mutational frequency detected by 2-kupl.

UBR4 contains 48 2-kupl mutations, more than any other gene. Fig. S4 shows read alignment at this gene for patient TCGA-EJ-7125 who carries the most UBR4 mutations (8/48 mutations). While seven of these mutations are absent in GDC portal variants, all can be visually validated as tumor-specific mutations as per the IGV display.

Besides recurrent mutations and indels, we found 20 genes with 43 recurrent structural variants predicted in at least two patients (suppl. Table S1). All these predicted variants can be supported by at least one read from the tumor library. Three recurrent structural variants map to prostate cancer genes *SH2B3, ATP10A* and *FOXA1* (Fig. 8). Variants in gene *ATP10A* and *SH2B3* have exactly the same junctions in at least two patients. As the three variants in gene *FOXA1* impact on the same exon, we grouped them as one same recurrent event despite not representing the exact same variation. All these recurrent structural variants are longer than 10bp. State-of-the-art procedures usually miss such variants at the mapping stage.

**Figure 8.**
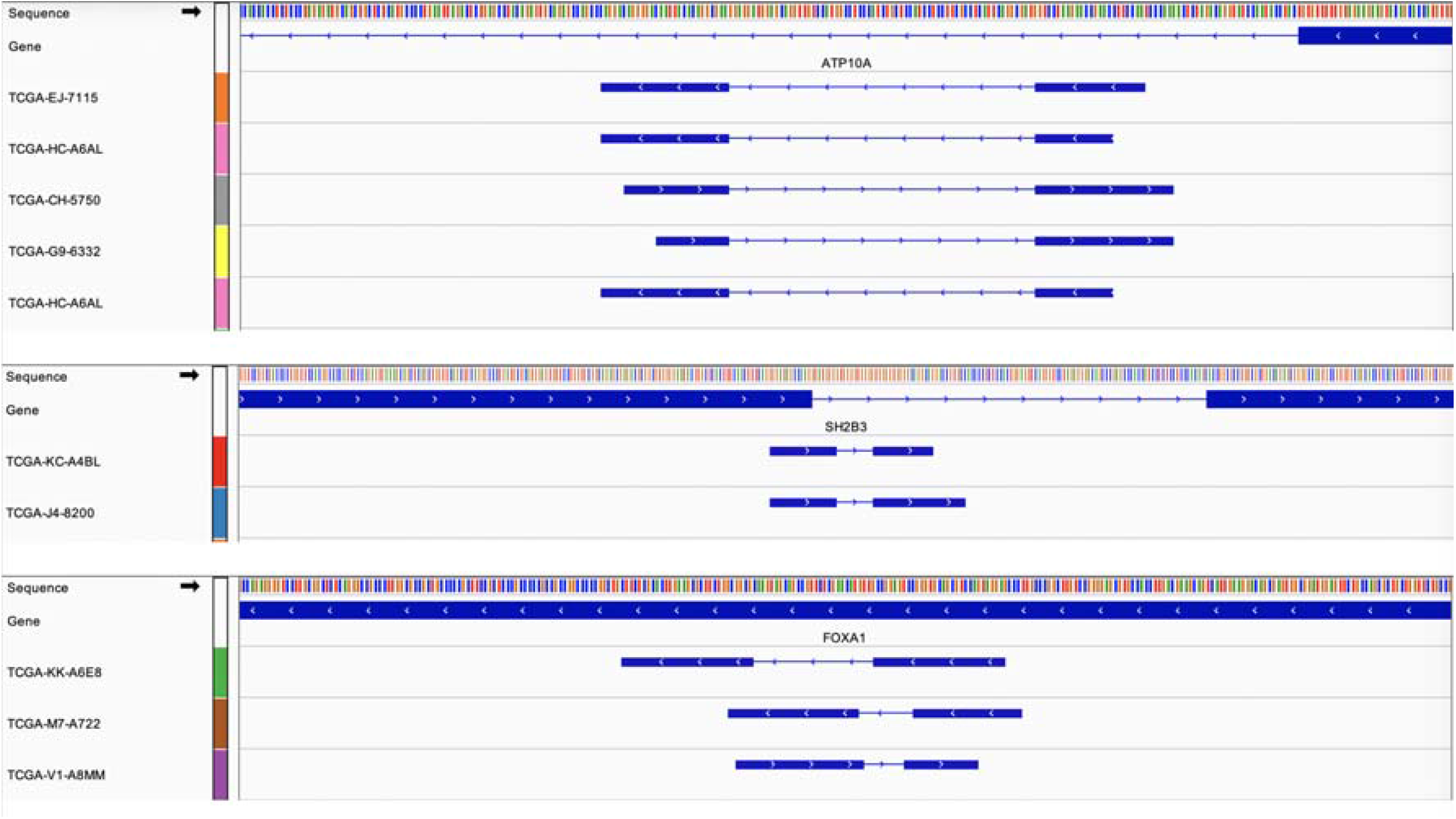
Recurrent structural variants mapping to three prostate cancer genes. In each track, lines represent the genome sequence (top), annotated genes, and variant contigs identified in different patients.

### Performance on bacterial WGS data

2-kupl can be applied to pairwise comparisons of DNA-seq datasets in any species. We present here an application to bacterial whole genome sequences. A frequent problem in bacterial genetics is identifying mutations in strains for which no reliable reference genome is available. We investigated the performance of 2-kupl on 21 DNA-seq datasets from *Pseudomonas aeruginosa*, in which 26 variants had been previously identified and confirmed by geneticists (see Methods).

About 141 variant contigs were predicted on average for each pair of WT/mutant strains, with an average running time of 10 minutes (Fig. 9A-B). Score ranking by 2-kupl and DiscoSNP++ allowed a clear separation of TP from FP (Fig. 9C-D). True positive calls were ranked first in 19 out of 19 mutant samples by 2-kupl and in 16 out of 16 samples by DiscoSNP++. Compared with Phred scores used in 2-kupl, DiscoSNP++ scales the rank scores from zero to one and thus the true positive variants are more concentrated.

**Figure 9.**
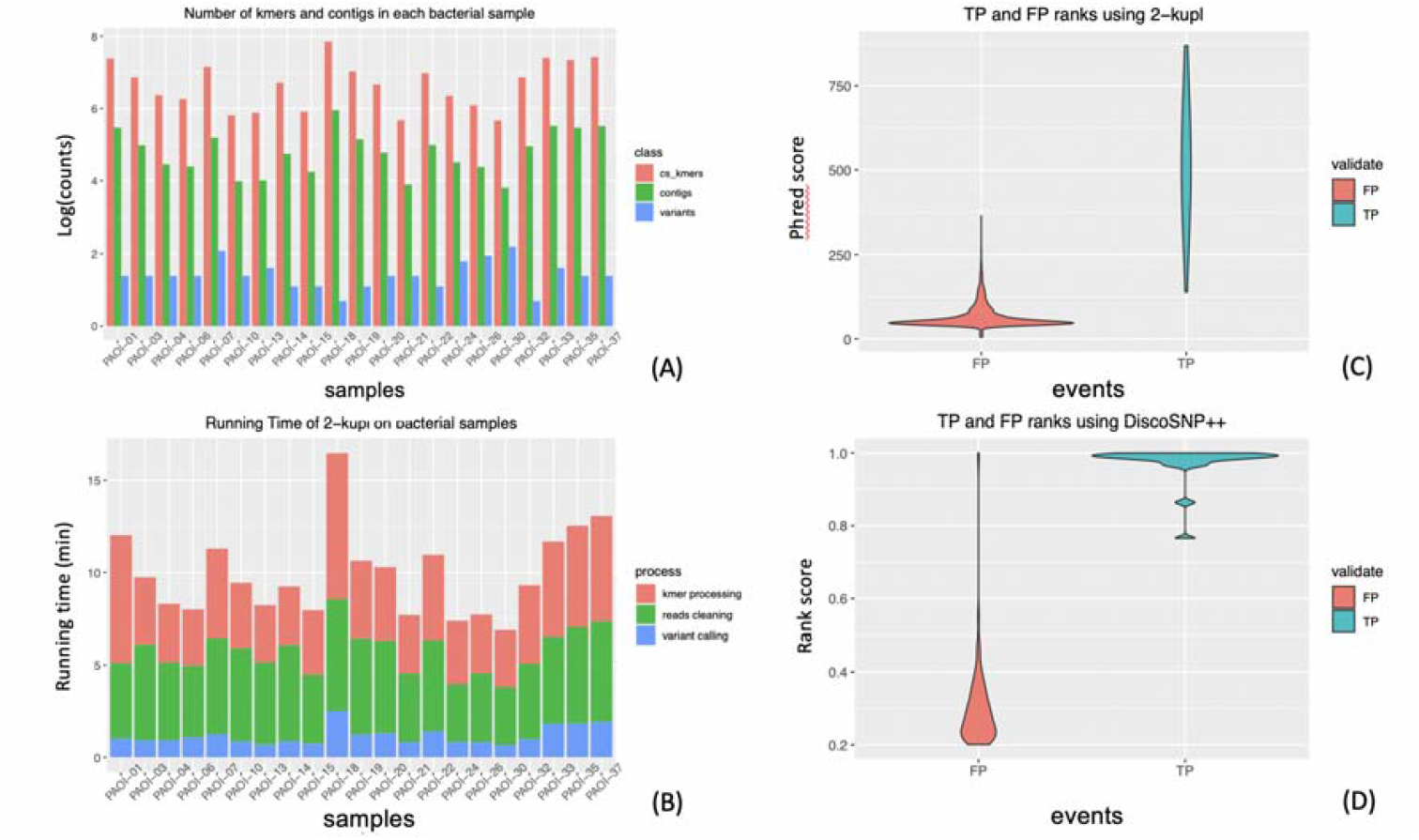
Performance of 2-kupl on bacterial DNA-seq datasets. (A) Number of cs_kmers, contigs and variants are shown for each bacterial sample. (B) Running time of 2-kupl on each sample is shown for different steps. (C) Distribution of Phred scores computed by 2-kupl in TP and FP events. (D) Distribution of DiscoSNP++ score ranks in TP and FP events.

2-kupl could recall all true positive variants, including SNVs and large deletions longer than 100 bp, while DiscoSNP++ missed three large deletions (555 bp, 213bp and 109 bp, suppl. Table S4). Meanwhile, DiscoSNP++ obtained 159 false positives vs. 45 for 2-kupl (Table 6). Therefore 2-kupl had the best recall and precision on this dataset, especially for large indels.

**Table 6.** comparison between 2-kupl and DiscoSNP++ on the bacteria DNA-seq data

|  | 2-kupl | DiscoSNP++ |
| --- | --- | --- |
| True Positive | 26 | 20 |
| False Positive | 45 | 159 |
| False Negative | 0 | 6 |
| Recall | 1 | 0.77 |
| FDR | 0.64 | 0.89 |
| Precision | 0.36 | 0.11 |
| F1 score | 0.52 | 0.20 |

## Discussion

Most variant detection protocols rely on reference genomes. However, even for species with a high-quality reference genome such as humans, depending on a reference is subject to limitations. Genomes contain large numbers of highly variable, repetitive or otherwise unmappable regions, which are unsolvable by short-read sequencing techniques. Hundreds of unsolved regions remain in telomeres and centromeres, also known as ‘dark matter’[56]. The X chromosome is the only complete human chromosome as of today[57]. Pathogenic variants within these unannotated regions are easily missed by mapping-based approaches due to low mapping quality, especially with low depth in whole-genome sequencing. Furthermore, the human genome varies across individuals and populations and a single reference genome does not account for this diversity[58].

2-kupl is able to detect variants, including mutations, indels and structural variants, without relying on a reference genome. Based on matched DNA-seq data, 2-kupl captures case-specific k-mers and counterpart k-mers (i.e. without the variation) into the same bucket. Sequence contigs harboring a local variation and its putative reference are inferred through the assembly of k-mers in each bucket.

To control artifacts induced by sequencing errors, 2-kupl takes both base quality and coverage into account. The general sequencing error rate in short-read NGS data is larger than 0.1%[59]. It is worth consuming computing resources and running time to remove these 0.1% artifacts because these sequencing errors result in large numbers of artifactual cs-kmers. To reduce the impact from low-quality bases, we combine Cutadapt and an ‘OverrideN’ function that flags low quality bases in the mid part of reads. This significantly reduces the number of cs-kmers and speeds up the computing procedure.

We compared the performance of 2-kupl with that of three competing methods in terms of running time, recall and precision. 2-kupl outperformed mapping-free methods DiscoSNP++ and Lancet in terms of recall and precision but did not reach the precision of the state-of-the-art alignment-based GATK-Mutect2 on human data.

DiscoSNP++ suffers from limitations of DBG data structures in regions with sequencing errors, genomic variants and repeats[18]. Efficient solutions searching for bubbles from such complicated structures are still under development. Furthermore, short contigs may be discarded within the post-process, cutting branches, for instance[60]. In our bacterial DNA-seq analysis, DiscoSNP++ missed some validated variants because of highly repetitive regions in the bacterial genome.

Lancet had a higher recall ratio than 2-kupl but also introduces more false positives. Furthermore, Lancet missed variants from repetitive regions and is not able to detect fusions from distant regions.

2-kupl had higher precision than DiscoSNP++ and Lancet with both human WES and bacterial DNA-seq data. Expectedly, 2-kupl did not outperform GATK-Mutect2 on WES data. First, GATK-Mutect2 uses a sophisticated Bayesian model to estimate a genotype’s likelihood given the observed sequence reads that cover the locus. When GATK-Mutect2 encounters a region showing signs of variation, it discards the existing mapping information and completely reassembles the reads in that region. This allows GATK-Mutect2 to be more accurate when calling regions that are traditionally difficult to call. Despite slightly fewer true positives, 2-kupl also detects fewer false positives than GATK-Mutect2. It is worth mentioning that 2-kupl has the lowest time complexity among the four methods.

By applying 2-kupl to the TCGA-PRAD patients, we were able to detect recurrent mutations and indels missed by the GDC portal’s GATK-Mutect2 pipeline. Reads in these regions have either low mapping qualities or multiple hits and were discarded in the GDC portal pipeline. Mapping-based methods all suffer from this issue and are powerless when faced with low complexity regions. 2-kupl identified recurrent mutations and recurrently mutated genes in high agreement with GATK-Mutect2. Mutated genes were enriched in PRAD-related genes, some of which specific to 2-kupl. As an example, we visually confirmed multiple 2-kupl-specific mutations in UBR4. Recurrent variants detected from the unmappable regions by 2-kupl provide insights into potential novel somatic variants even though the locus of origin of the contig sometimes cannot be determined.

Standard variant calling pipelines may miss mutations for multiple reasons: low allele frequencies, tumor contamination, ambiguities in short read alignment, inadequate sequencing depth, high GC content, sequencing errors and ambiguities in short read alignment. Different programs are affected by these factors to varying degrees. As a consequence, the mutations called by different pipelines are not consistent[61]. 2-kupl is not affected by some of these sources (GC content, alignment artifacts and mappability) and can detect a number of recurrent mutations (ie. potential driver events) that are not found by standard pipelines.

Several natural directions exist for extending 2-kupl. First, 2-kupl lacks sensitivity in detecting structural variants. All cs-kmers covering the junction are retained and extended to contigs. Unfortunately, neither the ct-kmers nor the reads are easily obtained when considering a hamming distance of one. A structural variation can be detected only if enough supporting reads are covering at least one side of the variation. Focusing on the cs-kmers regardless of ct-kmers could address this problem but at the cost of more false positives. A second limitation occurs when control samples are contaminated with tumor cells, which is relatively frequent in tissue biopsies. To address this problem, 2-kupl includes a parameter representing a k-mer count threshold in the control sample. However, a fixed contamination threshold may introduce unwanted non-specific variants. Future works should evaluate probabilistic approaches to address this issue.

## Declarations

### Ethics approval and consent to participate

Not applicable

### Consent for publication

Not applicable.

## Availability of data and materials

2-kupl is open source under MIT license and available at GitHub https://github.com/yunfengwang0317/2-kupl

## Competing interests

The authors declare that they have no competing interests.

## Funding

This work was funded in part by Agence Nationale de la Recherche grant ANR-18-CE45-0020 and by a PhD studentship to YW by Annoroad Technology, Beijing.

## Authors’ contributions

YW and DG designed the software and benchmarking procedures, YW developed the software, ran all analyzes and analyzed results, HX contributed to the software, YD reviewed and commented on the draft paper, CP provided bacterial genome data and analyzed results, YW and DG wrote the manuscript.

## Acknowledgements

Not applicable

## Notes

### Competing Interest Statement

The authors have declared no competing interest.

## Reference

1. Li MM, Datto M, Duncavage EJ, Kulkarni S, Lindeman NI, Roy S, et al. Standards and Guidelines for the Interpretation and Reporting of Sequence Variants in Cancer. J Mol Diagnostics. 2017.

2. MacArthur DG, Manolio TA, Dimmock DP, Rehm HL, Shendure J, Abecasis GR, et al. Guidelines for investigating causality of sequence variants in human disease. Nature. 2014.

3. Jiang T, Shi T, Zhang H, Hu J, Song Y, Wei J, et al. Tumor neoantigens: From basic research to clinical applications. Journal of Hematology and Oncology. 2019.

4. Shiloach J, Reshamwala S, Noronha SB, Negrete A. Analyzing metabolic variations in different bacterial strains, historical perspectives and current trends - example E. coli. Current Opinion in Biotechnology. 2010.

5. Hurles ME, Dermitzakis ET, Tyler-Smith C. The functional impact of structural variation in humans. Trends in Genetics. 2008.

6. Li H, Durbin R. Fast and accurate short read alignment with Burrows-Wheeler transform. Bioinformatics. 2009.

7. Tools VD. MuTect2. GATK Man. 2017.

8. Koboldt DC, Zhang Q, Larson DE, Shen D, McLellan MD, Lin L, et al. VarScan 2: Somatic mutation and copy number alteration discovery in cancer by exome sequencing. Genome Res. 2012.

9. Larson DE, Harris CC, Chen K, Koboldt DC, Abbott TE, Dooling DJ, et al. Somaticsniper: Identification of somatic point mutations in whole genome sequencing data. Bioinformatics. 2012.

10. Fan Y, Xi L, Hughes DST, Zhang J, Zhang J, Futreal PA, et al. MuSE: accounting for tumor heterogeneity using a sample-specific error model improves sensitivity and specificity in mutation calling from sequencing data. Genome Biol. 2016.

11. Li H, Ruan J, Durbin R. Mapping short DNA sequencing reads and calling variants using mapping quality scores. Genome Res. 2008.

12. Loeffler C, Karlsberg A, Martin LS, Eskin E, Koslicki D, Mangul S. Improving the usability and comprehensiveness of microbial databases. BMC Biol. 2020.

13. Audano PA, Ravishankar S, Vannberg FO. Mapping-free variant calling using haplotype reconstruction from k-mer frequencies. Bioinformatics. 2018.

14. Compeau PEC, Pevzner PA, Tesler G. How to apply de Bruijn graphs to genome assembly. Nat Biotechnol. 2011.

15. Uricaru R, Rizk G, Lacroix V, Quillery E, Plantard O, Chikhi R, et al. Reference-free detection of isolated SNPs. Nucleic Acids Res. 2015.

16. Narzisi G, Corvelo A, Arora K, Bergmann E, Shah M, Musunuri R, et al. Lancet: genome-wide somatic variant calling using localized colored DeBruijn graphs. Genome-wide Somat Var Call using localized Color Bruijn graphs. 2017.

17. Iqbal Z, Caccamo M, Turner I, Flicek P, McVean G. De novo assembly and genotyping of variants using colored de Bruijn graphs. Nat Genet. 2012.

18. Heydari M, Miclotte G, Van De Peer Y, Fostier J. Illumina error correction near highly repetitive DNA regions improves de novo genome assembly. BMC Bioinformatics. 2019.

19. Tomczak K, Czerwińska P, Wiznerowicz M. The Cancer Genome Atlas (TCGA): An immeasurable source of knowledge. Wspolczesna Onkologia. 2015.

20. Marçais G, Kingsford C. A fast, lock-free approach for efficient parallel counting of occurrences of k-mers. Bioinformatics. 2011.

21. Wu TD, Reeder J, Lawrence M, Becker G, Brauer MJ. GMAP and GSNAP for genomic sequence alignment: Enhancements to speed, accuracy, and functionality. In: Methods in Molecular Biology. 2016.

22. Chen C, Khaleel SS, Huang H, Wu CH. Software for pre-processing Illumina next-generation sequencing short read sequences. Source Code Biol Med. 2014.

23. Audoux J, Philippe N, Chikhi R, Salson M, Gallopin M, Gabriel M, et al. DE-kupl: Exhaustive capture of biological variation in RNA-seq data through k-mer decomposition. Genome Biol. 2017.

24. Pereira F, Azevedo F, Carvalho Â, Ribeiro GF, Budde MW, Johansson B. Pydna: A simulation and documentation tool for DNA assembly strategies using python. BMC Bioinformatics. 2015.

25. Bushnell B. BBMap. https://sourceforge.net/projects/bbmap/. 2015.

26. Huang X, Madan A. CAP3: A DNA sequence assembly program. Genome Res. 1999.

27. Wang Q, Shashikant CS, Jensen M, Altman NS, Girirajan S. Novel metrics to measure coverage in whole exome sequencing datasets reveal local and global non-uniformity. Sci Rep. 2017.

28. Danecek P, Auton A, Abecasis G, Albers CA, Banks E, DePristo MA, et al. The variant call format and VCFtools. Bioinformatics. 2011.

29. Cock PJA, Antao T, Chang JT, Chapman BA, Cox CJ, Dalke A, et al. Biopython: Freely available Python tools for computational molecular biology and bioinformatics. Bioinformatics. 2009.

30. McGinnis S, Madden TL. BLAST: At the core of a powerful and diverse set of sequence analysis tools. Nucleic Acids Res. 2004.

31. Li Z, Chen Y, Mu D, Yuan J, Shi Y, Zhang H, et al. Comparison of the two major classes of assembly algorithms: Overlap-layout-consensus and de-bruijn-graph. Brief Funct Genomics. 2012.

32. Robinson JT, Thorvaldsdóttir H, Winckler W, Guttman M, Lander ES, Getz G, et al. Integrative genomics viewer. Nature Biotechnology. 2011.

33. Meng J, Chen YPP. A database of simulated tumor genomes towards accurate detection of somatic small variants in cancer. PLoS One. 2018.

34. Zook JM, Catoe D, McDaniel J, Vang L, Spies N, Sidow A, et al. Extensive sequencing of seven human genomes to characterize benchmark reference materials. Sci Data. 2016.

35. Ewing AD, Houlahan KE, Hu Y, Ellrott K, Caloian C, Yamaguchi TN, et al. Combining tumor genome simulation with crowdsourcing to benchmark somatic single-nucleotide-variant detection. Nat Methods. 2015.

36. Abeshouse A, Ahn J, Akbani R, Ally A, Amin S, Andry CD, et al. The Molecular Taxonomy of Primary Prostate Cancer. Cell. 2015.

37. Tryka KA, Hao L, Sturcke A, Jin Y, Wang ZY, Ziyabari L, et al. NCBI’s database of genotypes and phenotypes: DbGaP. Nucleic Acids Res. 2014.

38. Mayakonda A, Lin DC, Assenov Y, Plass C, Koeffler HP. Maftools: Efficient and comprehensive analysis of somatic variants in cancer. Genome Res. 2018.

39. Landrum MJ, Lee JM, Benson M, Brown GR, Chao C, Chitipiralla S, et al. ClinVar: Improving access to variant interpretations and supporting evidence. Nucleic Acids Res. 2018.

40. Tate JG, Bamford S, Jubb HC, Sondka Z, Beare DM, Bindal N, et al. COSMIC: The Catalogue Of Somatic Mutations In Cancer. Nucleic Acids Res. 2019.

41. Pletscher-Frankild S, Pallejà A, Tsafou K, Binder JX, Jensen LJ. DISEASES: Text mining and data integration of disease-gene associations. Methods. 2015.

42. Kanehisa M, Furumichi M, Tanabe M, Sato Y, Morishima K. KEGG: New perspectives on genomes, pathways, diseases and drugs. Nucleic Acids Res. 2017.

43. Hamosh A, Scott AF, Amberger JS, Bocchini CA, McKusick VA. Online Mendelian Inheritance in Man (OMIM), a knowledgebase of human genes and genetic disorders. Nucleic Acids Res. 2005.

44. Ramos EM, Hoffman D, Junkins HA, Maglott D, Phan L, Sherry ST, et al. Phenotype-genotype integrator (PheGenI): Synthesizing genome-wide association study (GWAS) data with existing genomic resources. Eur J Hum Genet. 2014.

45. Martincorena I, Raine KM, Gerstung M, Dawson KJ, Haase K, Van Loo P, et al. Universal Patterns of Selection in Cancer and Somatic Tissues. Cell. 2017.

46. Armenia J, Wankowicz SAM, Liu D, Gao J, Kundra R, Reznik E, et al. The long tail of oncogenic drivers in prostate cancer. Nat Genet. 2018.

47. Latino L, Midoux C, Hauck Y, Vergnaud G, Pourcel C. Pseudolysogeny and sequential mutations build multiresistance to virulent bacteriophages in pseudomonas aeruginosa. Microbiol (United Kingdom). 2016.

48. de Koning APJ, Gu W, Castoe TA, Batzer MA, Pollock DD. Repetitive elements may comprise over Two-Thirds of the human genome. PLoS Genet. 2011.

49. Amemiya HM, Kundaje A, Boyle AP. The ENCODE Blacklist: Identification of Problematic Regions of the Genome. Sci Rep. 2019.

50. Pon JR, Marra MA. Driver and passenger mutations in cancer. Annu Rev Pathol Mech Dis. 2015.

51. Greenman C, Stephens P, Smith R, Dalgliesh GL, Hunter C, Bignell G, et al. Patterns of somatic mutation in human cancer genomes. Nature. 2007.

52. Goncearenco A, Rager SL, Li M, Sang QX, Rogozin IB, Panchenko AR. Exploring background mutational processes to decipher cancer genetic heterogeneity. Nucleic Acids Res. 2017.

53. Barbieri CE, Baca SC, Lawrence MS, Demichelis F, Blattner M, Theurillat JP, et al. Exome sequencing identifies recurrent SPOP, FOXA1 and MED12 mutations in prostate cancer. Nat Genet. 2012.

54. Fraser M, Sabelnykova VY, Yamaguchi TN, Heisler LE, Livingstone J, Huang V, et al. Genomic hallmarks of localized, non-indolent prostate cancer. Nature. 2017.

55. Li J, Drubay D, Michiels S, Gautheret D. Mining the coding and non-coding genome for cancer drivers. Cancer Letters. 2015.

56. Blaxter M. Revealing the dark matter of the genome. Science. 2010.

57. Miga KH, Koren S, Rhie A, Vollger MR, Gershman A, Bzikadze A, et al. Telomere-to-telomere assembly of a complete human X chromosome. Nature. 2020.

58. Sherman RM, Forman J, Antonescu V, Puiu D, Daya M, Rafaels N, et al. Assembly of a pan-genome from deep sequencing of 910 humans of African descent. Nature Genetics. 2019.

59. Ma X, Shao Y, Tian L, Flasch DA, Mulder HL, Edmonson MN, et al. Analysis of error profiles in deep next-generation sequencing data. Genome Biol. 2019.

60. Medvedev P, Pham S, Chaisson M, Tesler G, Pevzner P. Paired de Bruijn graphs: A novel approach for incorporating mate pair information into genome assemblers. J Comput Biol. 2011.

61. Hwang S, Kim E, Lee I, Marcotte EM. Systematic comparison of variant calling pipelines using gold standard personal exome variants. Sci Rep. 2015.

