## Supplementary figures and images for "2-kupl: mapping-free variant detection from DNA-seq data of matched samples"

### Supplemental Figure S1

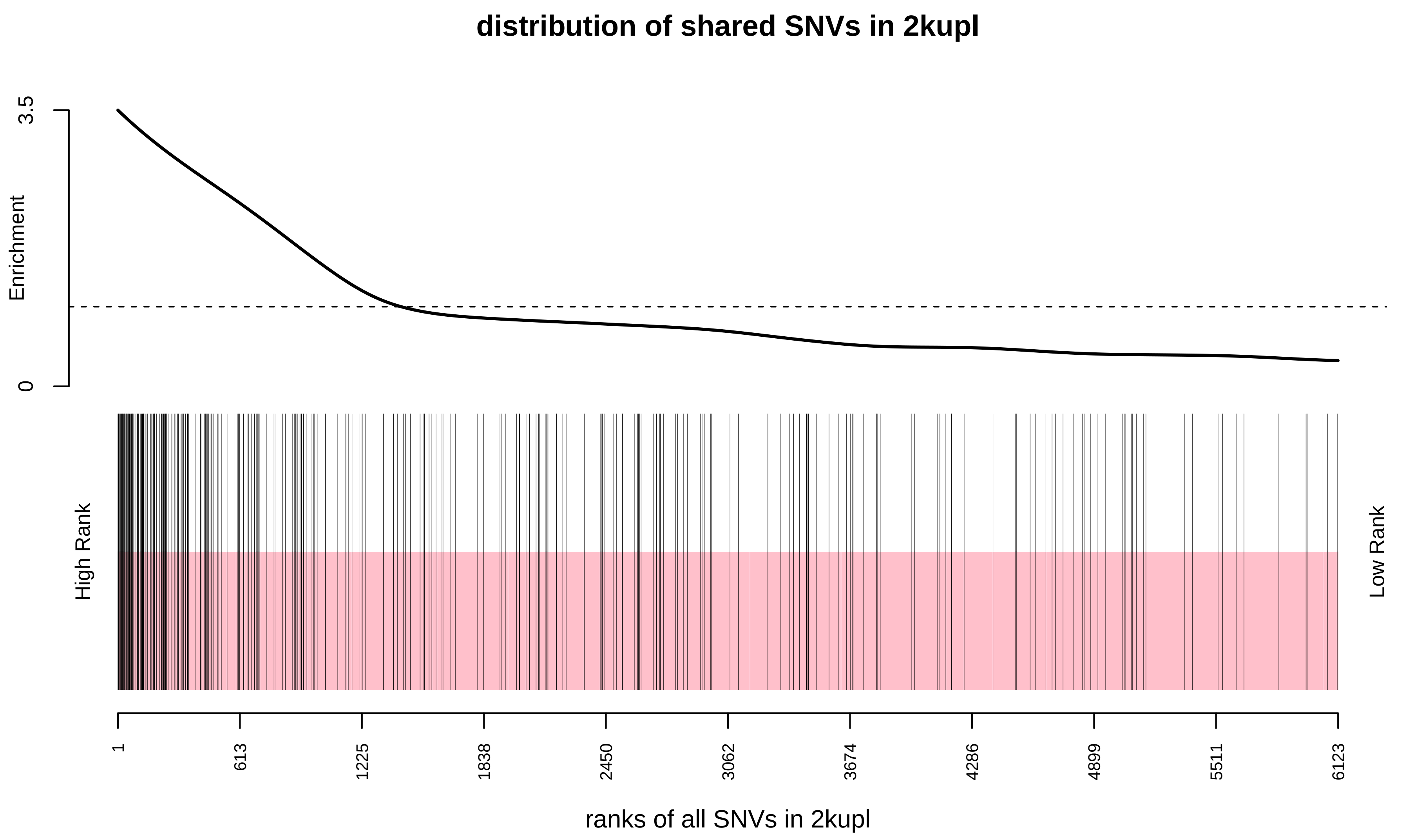

### Supplemental Figure S2

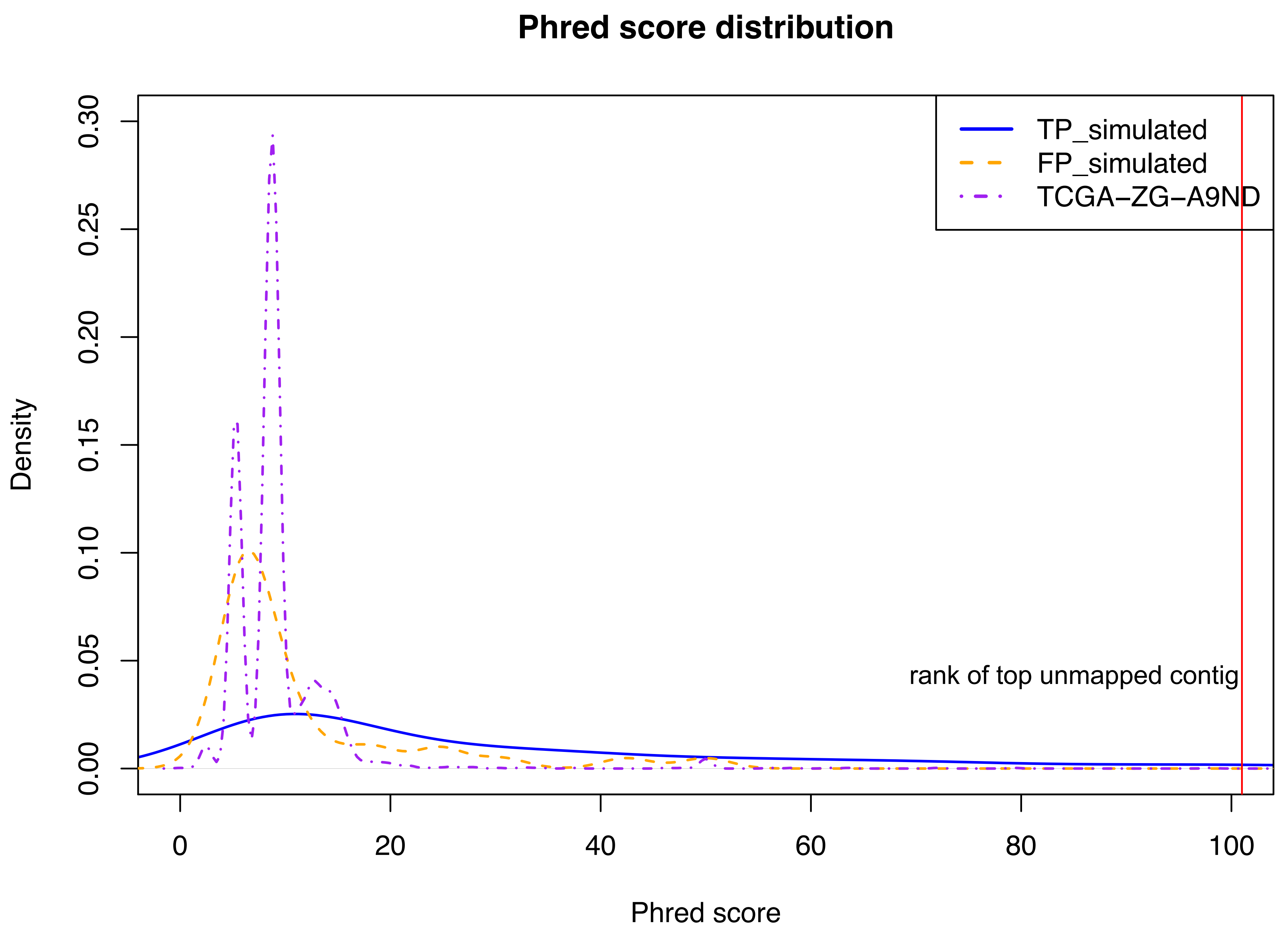

### Supplemental Figure S4

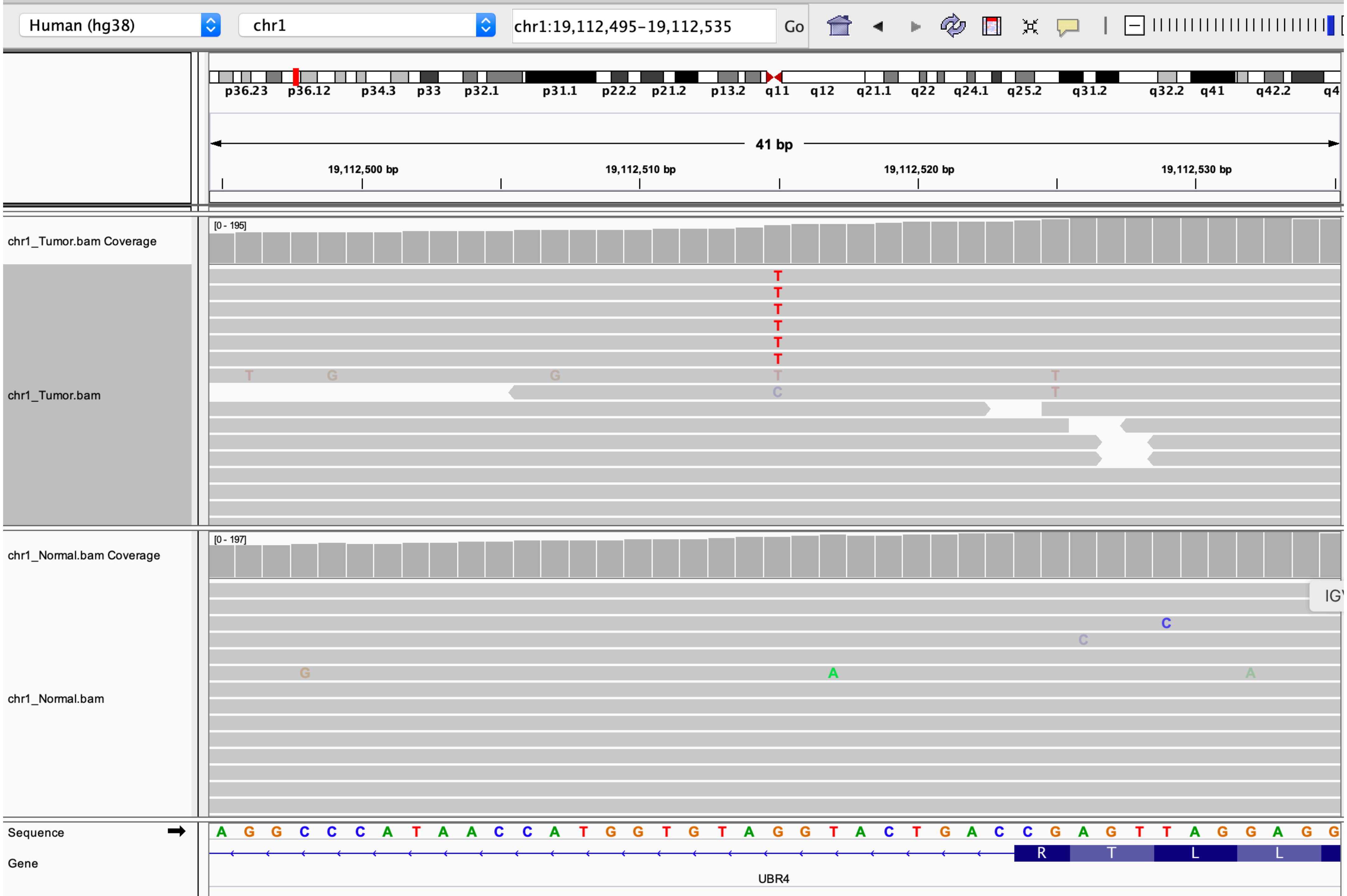

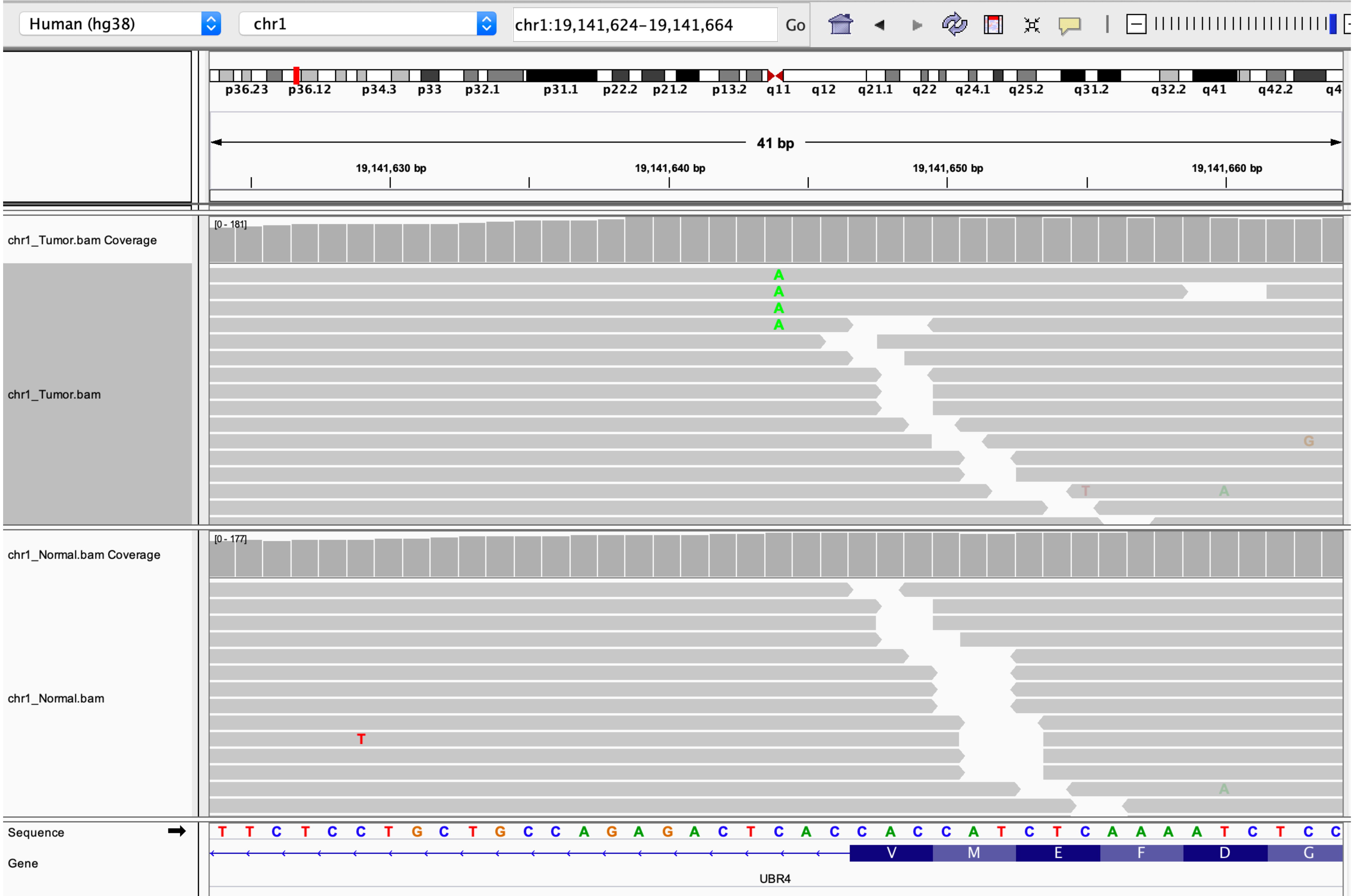

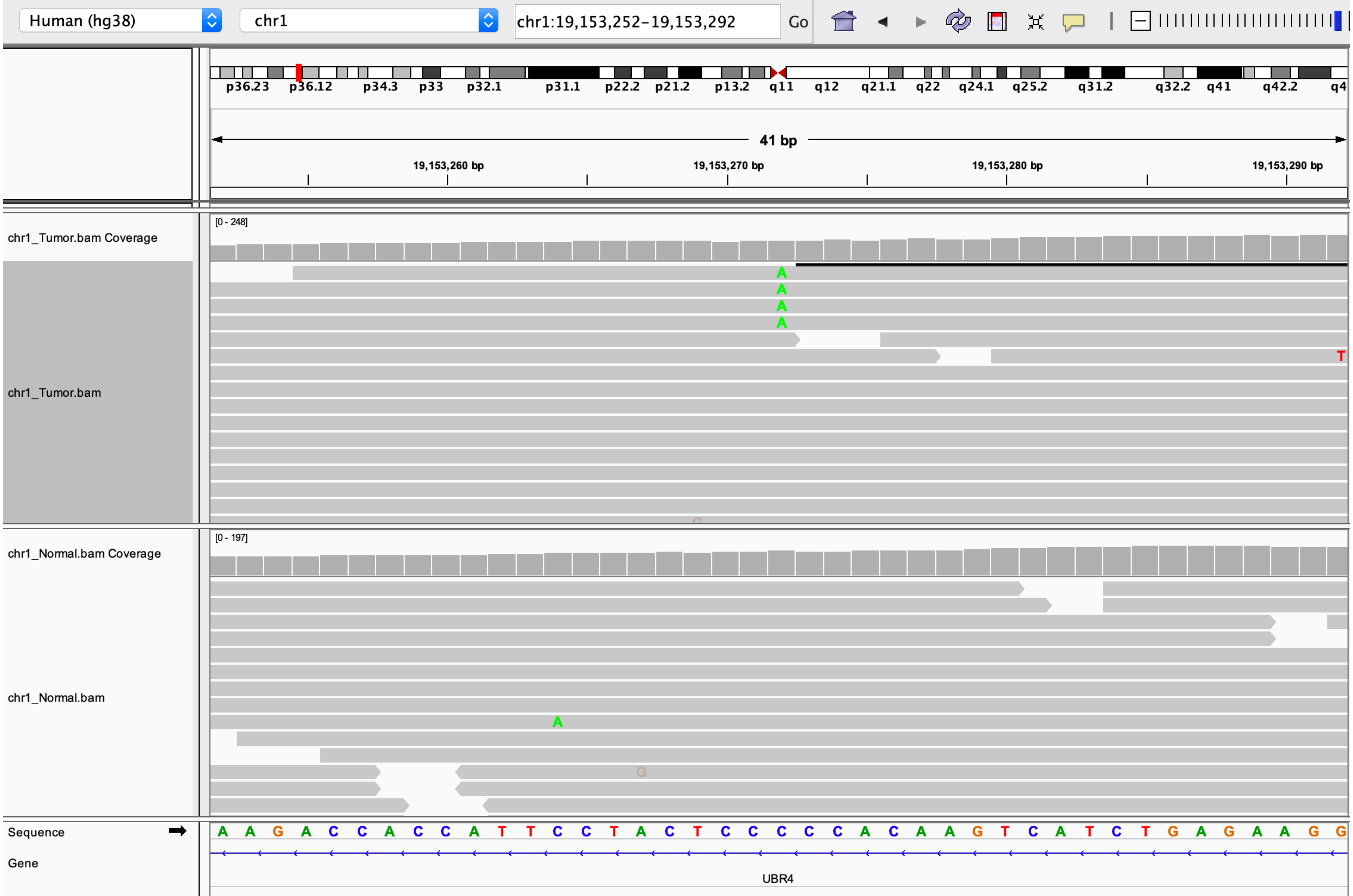

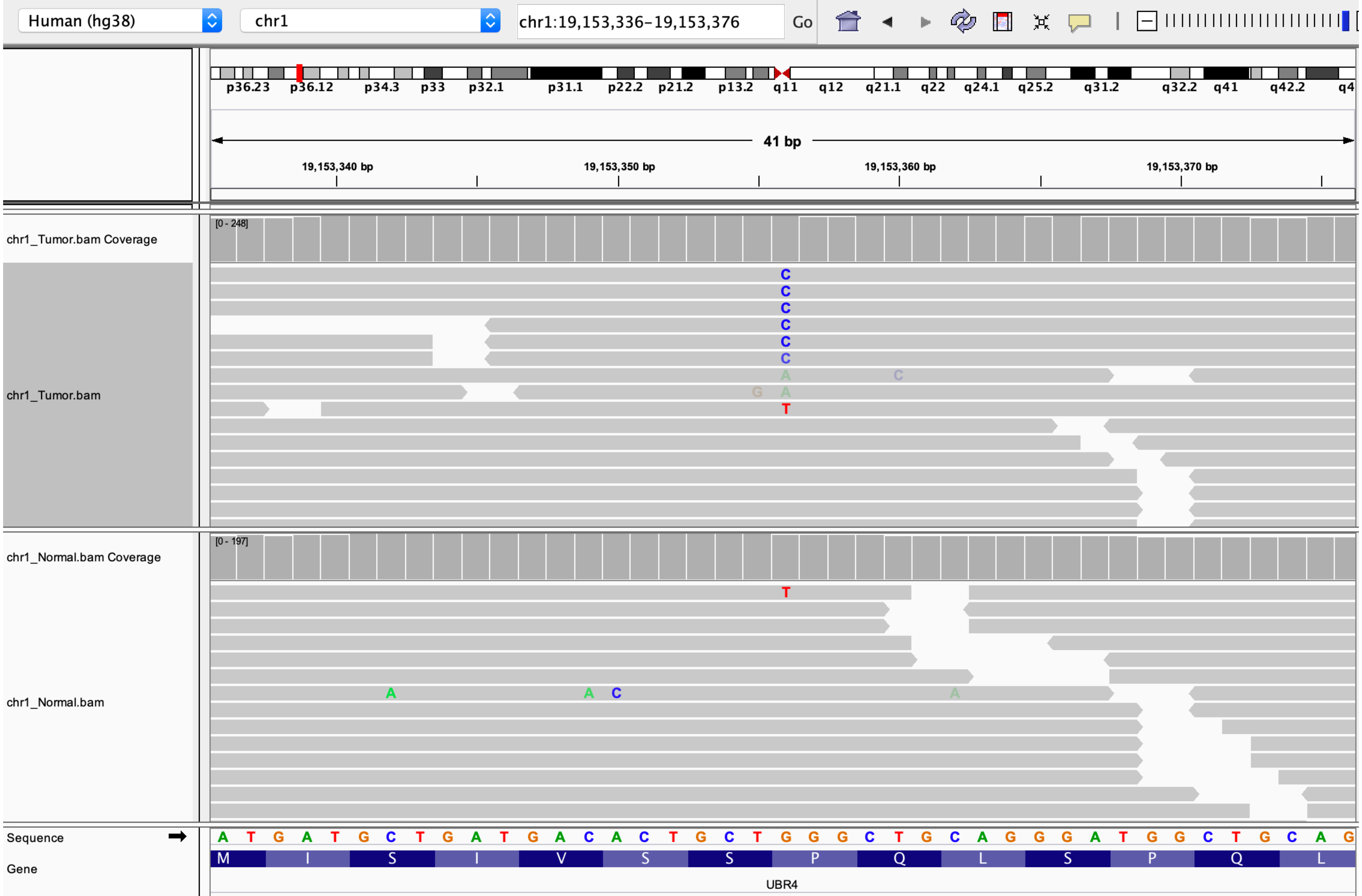

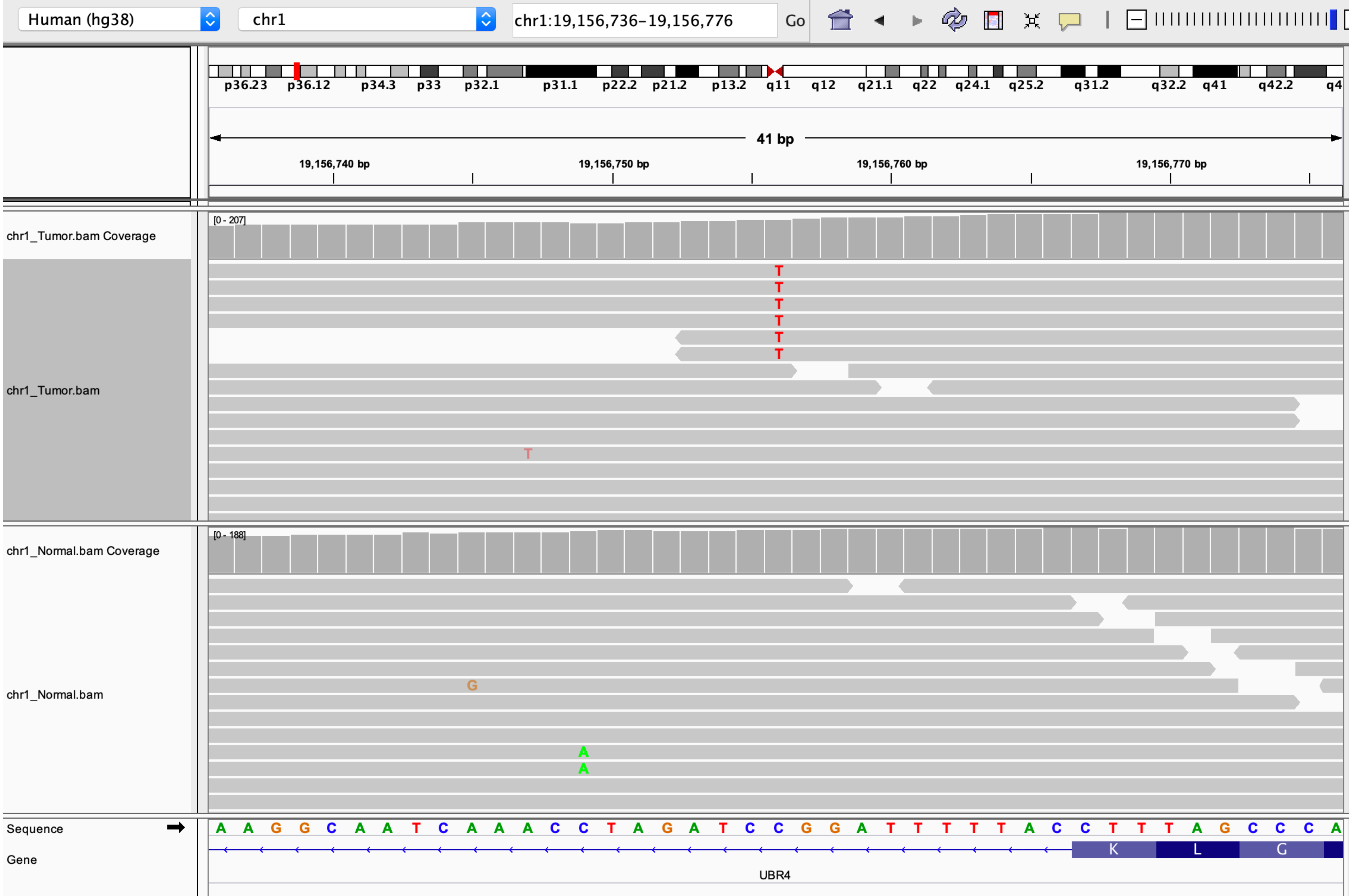

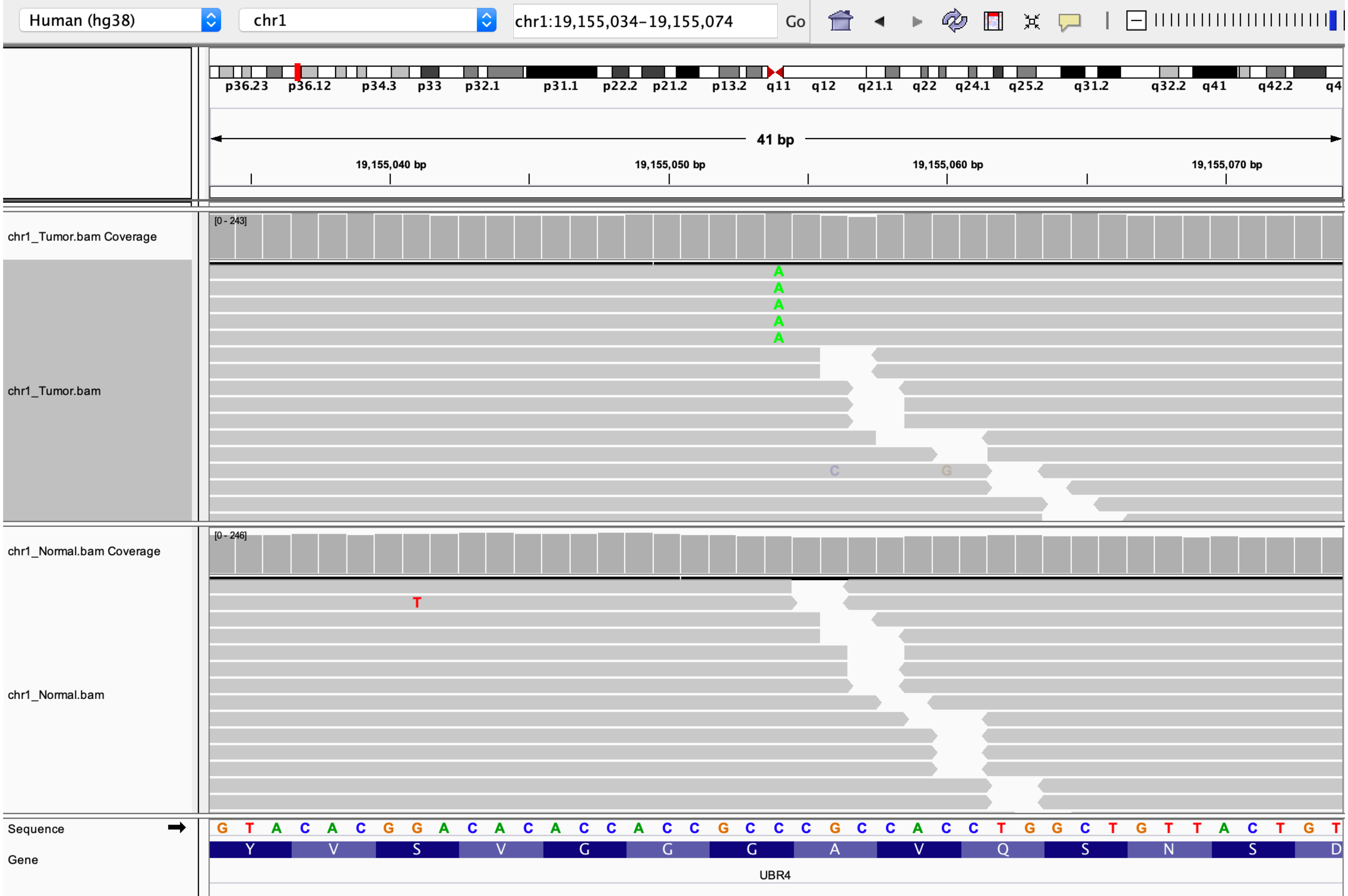

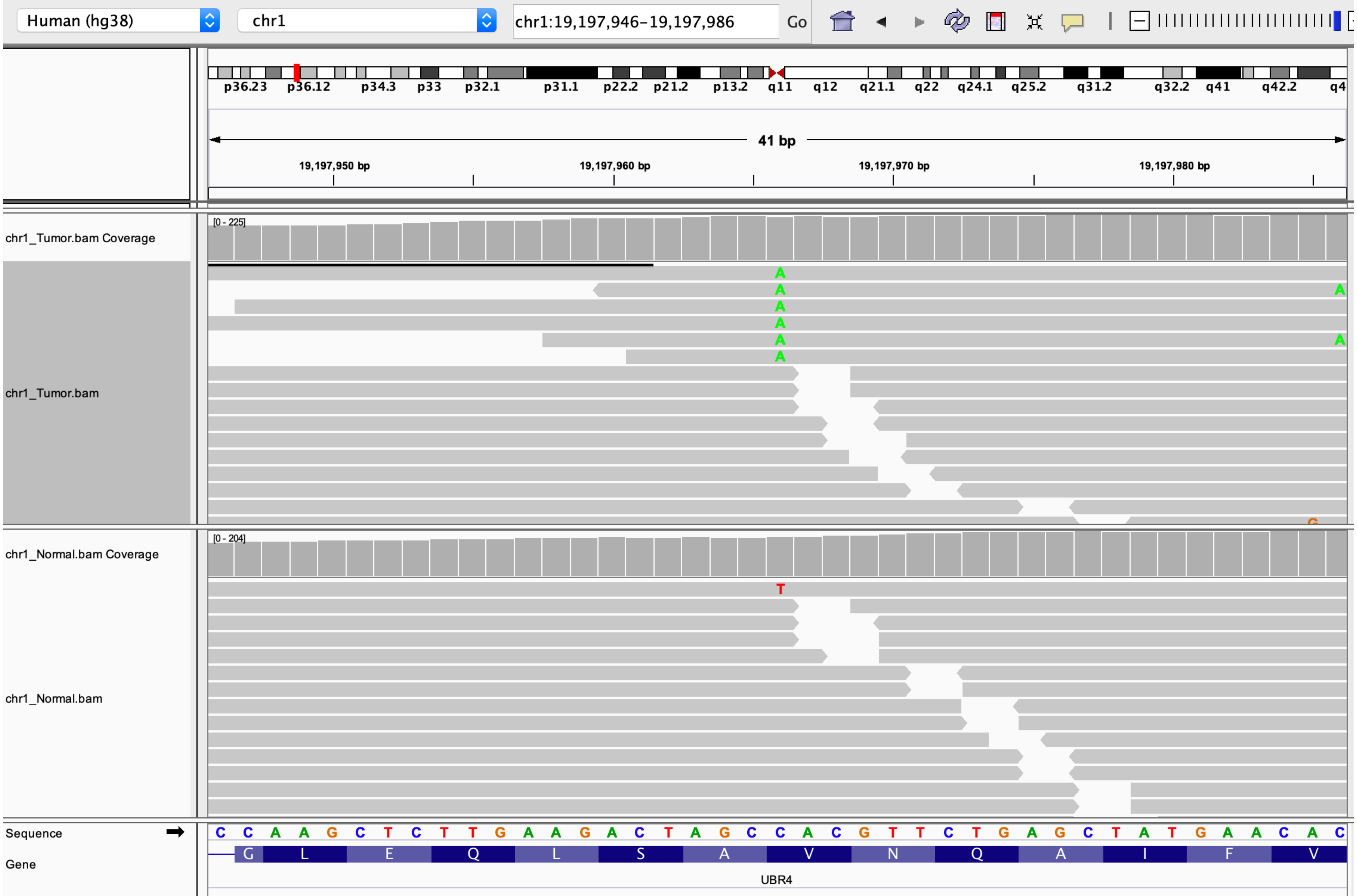
