## Supplemental Figure S3 for "2-kupl: mapping-free variant detection from DNA-seq data of matched samples"

```
GGGAGGGGTACTGTCACTCTGTTGCAGTAACAAGTTGCAAAATCTTCAGGCTGCAGGCTGCT
||||||||||||||||||||||||||||||.|||||||||||||||||||||||||||||||
GGGAGGGGTAATAAGTTGCAAAATCTTCAGGCTGCAGGCTGCT
Score=302.5   47      112     0       88
```
